# The mutational landscape of human somatic and germline cells

**DOI:** 10.1101/2020.11.25.398172

**Authors:** Luiza Moore, Alex Cagan, Tim H.H. Coorens, Matthew D.C. Neville, Rashesh Sanghvi, Mathijs A. Sanders, Thomas R.W. Oliver, Daniel Leongamornlert, Peter Ellis, Ayesha Noorani, Thomas J Mitchell, Timothy M. Butler, Yvette Hooks, Anne Y. Warren, Mette Jorgensen, Kevin J. Dawson, Andrew Menzies, Laura O’Neill, Calli Latimer, Mabel Teng, Ruben van Boxtel, Christine A. Iacobuzio-Donahue, Inigo Martincorena, Rakesh Heer, Peter J. Campbell, Rebecca C. Fitzgerald, Michael R. Stratton, Raheleh Rahbari

**Author notes:** joint first authors (Luiza Moore, Alex Cagan, Tim H.H. Coorens). Correspondence to (R.R.) and (M.R.S.).

## Abstract

During the course of a lifetime normal human cells accumulate mutations. Here, using multiple samples from the same individuals we compared the mutational landscape in 29 anatomical structures from soma and the germline. Two ubiquitous mutational signatures, SBS1 and SBS5/40, accounted for the majority of acquired mutations in most cell types but their absolute and relative contributions varied substantially. SBS18, potentially reflecting oxidative damage, and several additional signatures attributed to exogenous and endogenous exposures contributed mutations to subsets of cell types. The mutation rate was lowest in spermatogonia, the stem cell from which sperm are generated and from which most genetic variation in the human population is thought to originate. This was due to low rates of ubiquitous mutation processes and may be partially attributable to a low cell division rate of basal spermatogonia. The results provide important insights into how mutational processes affect the soma and germline.

## Introduction

Studying mutations arising in normal human cells during the lifetime of an individual provides insight into the development, maintenance and structure of normal tissues (Coorens et al. 2020), the mutational processes that have been operative, and the role of selection in shaping cell populations. It can elucidate how each of these are altered by, or contribute to, cancer, other diseases, and ageing.

Characterising such mutations has been technically challenging, as normal cell populations consist of myriad small clones, with the mutations differing between clones. Recently, several approaches have been developed to identify mutations in normal tissues. These include i) sequencing cell populations expanded from single cells by *in vitro* culture^1–5^; ii) sequencing small pieces of tissue to high depth to detect mutations present in small clonal populations^6–8^; iii) sequencing individual microscopically visible tissue structures derived from single cells^9–11^, iv) sequencing DNA from single cells^12,13^; v) duplex DNA sequencing, in which information from both DNA strands is used to detect rare mutations^14–16^. Each has its own strengths and limitations.

The body is composed of somatic and germline cells. Mutations acquired in somatic cells during a lifespan, and their consequences, are restricted to the individual in whom they occur. A subset of somatic cell types have been investigated in depth and differences between them in clonal structure, mutation rates and processes, and frequency of drivers reported^2,9–11^. However, most somatic cell types remain to be investigated. Variation between individuals has also been observed, indicating that genetic, environmental and lifestyle factors may influence patterns of somatic mutation.

By contrast, mutations in the germline can be transmitted to the next generation, making them the raw material of species evolution and the cause of hereditary diseases. Understanding of mutagenesis in reproductive tissues that create germline mutations, their rates, and underlying processes has predominantly been inferred from nuclear-family (trio) studies, which have shown that ~80% of transmitted germline mutations arise in the paternal germline^17–19^. There have been inferences that somatic and germline mutation rates differ^20,21^ but there has been no direct comparison of germline and somatic mutation rates and processes and little is known about mutagenesis in the cell lineage leading to sperm.

Here, we employed a design in which multiple cell types from the same individuals were compared, thus controlling for interindividual genetic and environmental differences. We investigated the mutation landscape in a wide range of normal somatic tissues and compared it with the male germline.

## Results

### Microdissection of normal tissues

We laser microdissected 389 patches (200-1000 cells) of 29 distinct histological structures from three individuals, a 47-year-old male, a 54-year-old female and a 78-year-old male (**Supplementary Table 1,2**). DNA extracted from each patch was whole genome sequenced (WGS) to 27-fold median coverage using low-input library-construction methods (Ellis et al, 2020) (**Fig. 1a**). Overall, 484,678 single base substitutions (SBS), 8,388 small insertions and deletions (ID), 37 copy-number changes and 128 structural variants were identified (**Extended Data Fig. 1–4; Supplementary Table 2-6**).

**Figure 1.**
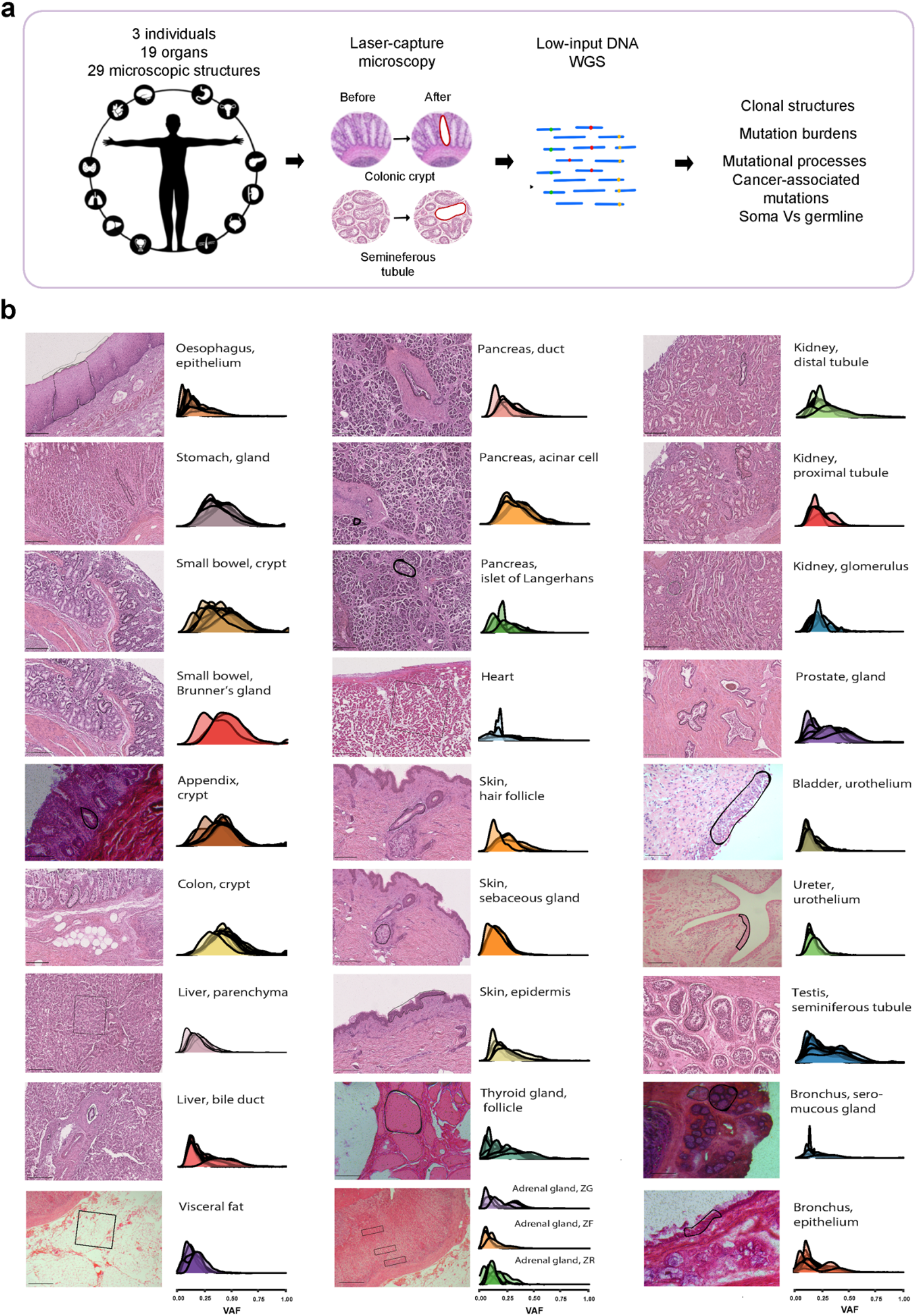
Summary of the experimental design and clonal dynamics across tissues. **a.** Each tissue biopsy was examined and specific populations of microscopic structures were laser-capture microdissected (LCM). The obtained cellular material was subjected to DNA extraction and whole genome sequencing (WGS) using a modified library-construction protocol. Mutations acquired during life were identified by comparison with WGS data extracted from macroscopic pieces of normal tissue from the same individuals. **b.** Histological sections and VAF distributions of somatic mutations from 389 patches across 29 microscopic structures.

### Clonal structures in normal tissues

Many cell types in the adult human body are renewed by stem cell division generating further proliferating and terminally differentiated cells^22^. For some cell types, descendants of a stem cell remain in close proximity forming localised clonal populations which may differ in size and shape between tissues and form microscopic anatomical structures, such as colorectal crypts and endometrial glands^9^. For others, descendants of a stem cell mingle with descendants of other stem cells to form polyclonal populations, for example in the blood^2^. The clonal architecture of a tissue thus reflects the way it is constituted and maintained.

Whether a cell population is derived from one or multiple stem cells can be inferred from the variant allele fraction (VAF) of acquired mutations. Microdissected biopsies from different tissues showed substantially different VAF distributions (**Fig. 1b**). Many discrete microanatomical glandular structures, including colorectal, small intestinal, appendiceal crypts, gastric glands and duodenal Brunner’s glands were usually monoclonal (median VAF 0.45, ranges 0.29-0.55). Prostatic glands and seminiferous tubules were often monoclonal (median VAF 0.31, ranges 0.16-0.46). Other cell types from ductal, tubular and some glandular structures, squamous epithelial sheets, and tissues without well-defined microstructure were infrequently monoclonal (median VAF 0.25, ranges 0.14-0.45). These were likely composed of mosaics of clones with microdissected biopsies usually including multiple clones, either because the clone size is smaller than the number of cells dissected or because, without microanatomical structures to guide microdissection, the tissue excised overlaps the boundaries between multiple clonal units. Patches from cardiac muscle, skeletal muscle and brain yielded very few mutations, consistent with these being primarily non-renewing in the adult and/or being composed of so many clones that none achieve the level of clonal dominance required for calling somatic mutations. The results therefore indicate that many tissues were composed of populations derived from single renewing cells and highlight the potential of DNA sequence-based approaches, coupled to microscopy, to further elucidate tissue architecture, cell lineages and cell dynamics.

Previous studies have revealed that driver mutations conferring selective advantage are present in normal tissues. Most cell types studied here showed none or a small number of driver mutations with hotspot canonical drivers in *BRAF* and *GNAS* in appendiceal crypts, *PTPN11* in seminiferous tubules, *FOXA1* in prostate, *KRAS* in small intestine and *TP53* in oesophagus as well as nine truncating mutations in eight recessive cancer genes across the sample set (**Supplementary Table 7**). Additionally, four chromosome-arm or focal losses, encompassing either *NOTCH1* and *TP53*, with damaging mutations on the other allele were observed in oesophagus (**Extended Data Fig. 5**).

### Mutation burdens and rates in normal tissues

Monoclonal populations permit making inferences about mutation burdens and rates that are not possible in polyclonal populations. Therefore, analysis of mutation burden and rate was limited to cell types in which most microdissected patches were dominated by a single clone and for which multiple monoclonal microbiopsies were available from more than one donor (**Methods**). With respect to somatic tissues, crypts of the large intestine, small intestine and appendix exhibited the highest mutation rates (52 SBS/year CI95% 48-54) with lower rates in gastric glands (25 SBS/year; CI95% 20-32), prostatic glands (19 SBS/year CI95% 17-22), pancreatic acini (15 SBS/year CI95% 8-23), and bile ductules (9 SBS/year CI95% 5-21) (**Extended Data Fig. 1,2**). Thus, differences in clonal dynamics exist between somatic cell types from the same individuals. These may be due to differences in mutation burden or in principal time to the most recent common ancestor (TMRCA).

We next explored the mutation burden in the male germline. Seminiferous tubules of the testis are lined by germinal epithelium composed of spermatogonial stem cells, a hierarchy of intermediate germ cells leading to sperm, and a small population of Sertoli cells supporting the germ cells. Thus, microdissections of seminiferous tubules are predominantly composed of germline cells. VAF distributions of mutations from these indicated that most were monoclonal, indicating that they derive from a single ancestral spermatogonium. Mutation burdens and rates were much lower (2.38/year CI95% 1.83-2.53) in seminiferous tubules than in the somatic cell types analysed. To further characterise these differences in mutation burden and rate between somatic and germline cells, 162 microbiopsies from the seminiferous tubules of an additional 11 men (ages 22-83 years, median 44 years) and 10 colorectal crypts from four of these donors (**Supplementary Table 1**) were whole genome sequenced. There was substantial variation in mutation burden in the seminiferous tubules from the 13 individuals ranging from 23 to 294 SBS (median 91 CI95% 80-108) and 1 to 17 indels (median 3 CI95% 2.21-3.36, **Extended Data Fig. 3**). This variation was mostly explained by a linear correlation with age and accumulation of ~2.6 SBS (mixed linear model, CI95% 2.1-3.1, P= 5.02 x 10^-7, R2= 0.71) and ~0.07 indels (mixed linear model, CI95% 0.02-0.13, P= 2.08 x 10^-2, R2= 0.31) indels per tubule per year. The SBS mutation rate in seminiferous tubules was ~27-fold lower than in colorectal crypts obtained from six of the individuals studied (**Fig. 2a**). The haploid SBS mutation rate in spermatogonial stem cells is consistent with the germline mutation rate of ~1.35 mutations per year in the paternal germline from “trio” studies^17,19,23^(**Fig. 4b; Methods**)

**Figure 2.**
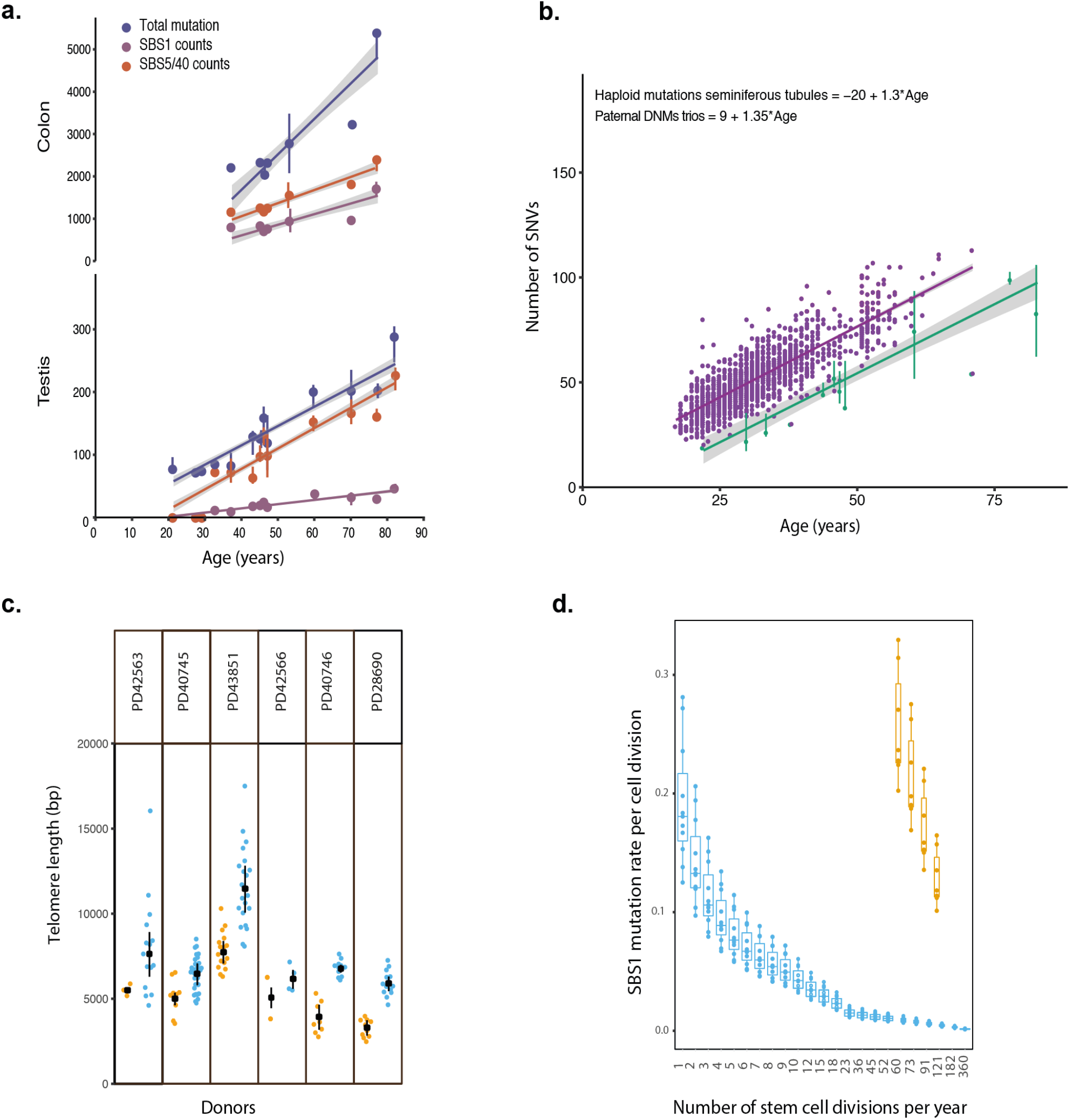
Mechanisms underlying the low germline mutation rate. **a.** Comparison of total (dark blue); SBS1 (orange); SBS5/40 (purple) mutation burden in the seminiferous tubules and matched colonic crypts of nine individuals **b**. Comparison of haploid mutation burden in seminiferous tubules (green) with paternal germline *de novo* mutations (purple) **c.** Relative telomere length in seminiferous tubules (blue) and colonic crypts (orange) from the same individuals. **d.** Model to compare SBS1 mutational burden in the spermatogonial stem cells (blue) and colonic crypts stem cells (orange). Ranges of stem cell divisions per year in spermatogonial stem cells were compared with colonic crypts (dividing every 2-5 days).

Telomere shortening is a hallmark of ageing. We measured the relative telomere length^24^ (**Methods**) across all cell types showing substantial variability between cell types and individuals (**Fig. 3c**). There was loss of 72bp per year in somatic cell types (p = 0.03289 likelihood-ratio test, **Fig. 3c** and **Extended Data Fig. 6**) but this decline with age was not observed in seminiferous tubules (P-value = 0.49). Indeed, seminiferous tubules predominantly showed longer telomere lengths than all somatic tissues sampled (median length ~7kb) with, for example, ~3kb (SD 968bp, range 1344-3489bp) longer telomeres than in colorectral crypts from the same individual (**Fig. 2c**).

**Figure 3.**
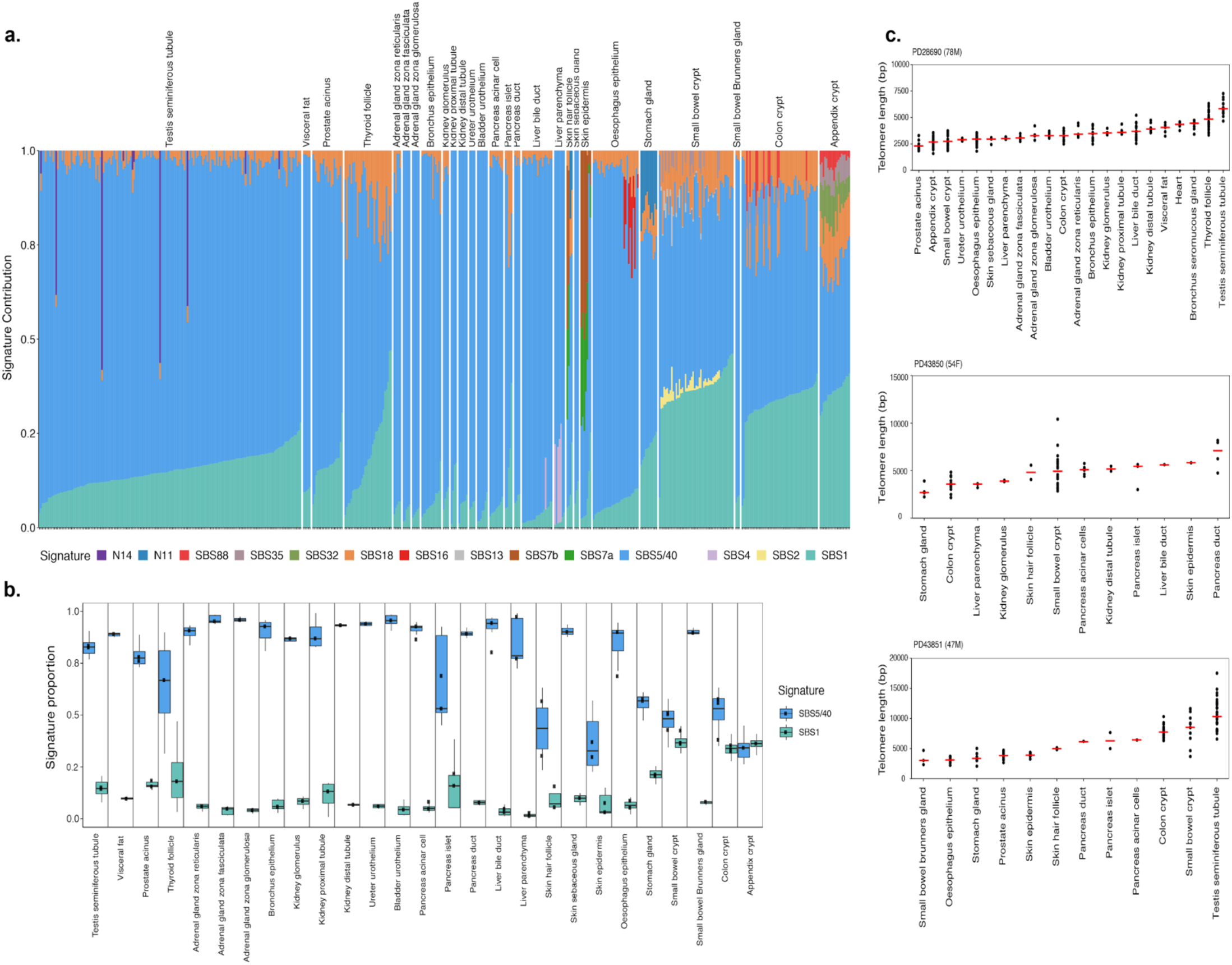
Mutational signatures in normal tissues and telomere lengths analysis. **a.** Mutational signatures and their relative contribution across normal tissues **b.** Variation in the relative contribution of SBS1 (green) and SBS5/40 (blue) across microscopic structures. **c.** Telomere length of microdissected patches per tissue per donor, estimated from WGS data.

### Mutational signatures in normal tissues

To explore the mutational processes operative in normal tissues we extracted mutational signatures (**Methods**) and estimated the mutation burden attributable to each signature (**Fig. 3a**). This analysis included all cell types and was not restricted to those with monoclonal populations. SBS1, which is likely due to deamination of 5-methylcytosine^25,26^, and SBS5, which is of unknown cause but thought to be a pervasive and relatively clock-like endogenous process^27^, were present in all the normal cell types which yielded mutations. Together they accounted for the large majority of mutations in all cell types (**Fig. 3b**). SBS1 and SBS5 are also ubiquitous among cancer types^28^ (**Extended Data Fig. 7**).

Other mutational signatures were restricted to subsets of normal cell types. SBS18, which may be due to oxidative damage^29^, was observed in many cell types (19/29) but generally constituted a higher proportion of mutations in large and small intestinal crypt stem cells compared to other cell types. SBS35, which is due to platinum chemotherapy agents^30^, was observed in sub-clonal populations of multiple cell types from an individual who received treatment two-months before death (**Supplementary Table 1**). SBS7, associated with UV-light exposure^31^, was found in all patches of skin epidermis and some hair follicles, but not in skin sebaceous glands. SBS88, due to a mutagenic product (known as colibactin) of a strain of *Escherichia coli* in the intestinal microbiome^9,32^, was found in a subset of colorectal and appendiceal crypts. SBS16, of unknown cause but previously associated with alcohol consumption in some cancers^33^, was observed in oesophagus. SBS2 and SBS13, likely due to APOBEC cytosine deaminases^34^, were found in a subset of small intestinal crypts. SBS4, which is associated with tobacco smoke exposure^35,36^, was found in a subset of liver parenchymal cells, as reported previously in both normal liver and liver cancer^10^. The reasons for the tissue specificities of SBS2, SBS13, SBS16 and SBS4 are unknown. Small contributions of SBS32, N11 and N14 were also found across multiple cell types and their significance is unknown. We cannot exclude the possibility that further signatures with low mutation burden or present in a few cell types are present. SBS40 and SBS5 are both flat and featureless signatures; it is difficult to accurately attribute mutation loads to each separately and we have, therefore, merged their burdens. Therefore, in normal somatic cell types (with the exception of skin) most mutations are caused by the mutational processes underlying SBS1 and SBS5/40. However, several other mutational processes make lesser contributions to particular cell types as a result of exogenous and endogenous exposures.

The relative proportions of SBS1 and SBS5/40 mutations differed between somatic cell types (**Fig. 3b**). For example, in colorectal crypts SBS1 accounted for 0.39(CI95% 0.375-0.435) of mutations and SBS5/40 for 0.61(CI95% 0.565-0.625), with similar proportions in the small intestine, whereas in prostatic glands the two signatures accounted for 0.17(CI95% 0.163-0.189) and 0.83(CI95% 0.811-0.837) of mutations respectively. Thus, although SBS1 and SBS5/40 both accumulate with age in normal somatic tissues^9^ “, there must be underlying mechanisms explaining the variation in the relative proportion of somatic mutations accounted for by SBS1 and SBS5/40. SBS1 mutations have previously been related to rates of cell division^27^ and it is plausible that the high SBS1 rates in colorectal stem cells compared to other cell types are due to higher rates of cell division. In the germline, SBS5/40 accounted for 0.85(CI95% 0.835-0.859) and SBS1 0.15(CI95% 0.141-0.165) of mutations in seminiferous tubules, with a small contribution from SBS18, a very similar landscape to that observed in *de novo* germline mutations inferred from trios^17^.

### Cellular mechanisms underlying the low germline mutational rate

The much lower mutation rate in seminiferous tubules compared to somatic cells is an important insight into maintenance of the human germline. The substantial difference between seminiferous tubules and colorectal crypts, in particular, provides an opportunity to explore mechanisms underlying differences in somatic and germline mutation rates.

In both seminiferous tubules and colorectal crypt stem cells, base substitution mutations were accumulated in a linear manner throughout life (**Fig. 2a**). Lower mutation rates of both SBS1 and SBS5/40 in seminiferous tubules accounted for most of the difference in mutation rate between seminiferous tubules and colonic crypts stem cells. SBS5/40 accounted for more of this difference than SBS1. However, the relative difference in SBS1 mutation rate (41-fold less in seminiferous tubules compared to colonic crypts) was much greater than the relative difference in SBS5/40 mutation rate (12-fold less in seminiferous tubules compared to colonic crypts). Thus, lower mutation rates of ubiquitous signatures accounted for most of the difference in mutation burden in seminiferous tubules compared to colonic crypts stem cells. The remainder could be explained by SBS18, which was present across colorectal crypt stem cells but rarely in seminiferous tubules where it contributed relatively few mutations, and the sporadic occurrence of the colibactin induced SBS88 in a minority of colorectal stem cells.

Cells may be particularly likely to acquire mutations during the DNA replication that takes place in S-phase of cell division. We therefore investigated whether the lower mutation rate in seminiferous tubules compared to colorectal crypts could be explained by a lower rate of spermatogonial cell division. Previously published data indicate that basal stem cells in human colorectal crypts divide every 2-4 days^37^ while the division rate for the basal spermatogonial stem cells remains more contentious, with estimates ranging from every 16 days^38^ to only a few times a year^39,40^. We modelled different rates of cell division for these two cell types assuming the same mutation rate per cell division and found that our results were most consistent with a scenario in which the basal spermatogonial stem cells are predominantly quiescent, dividing only a few times per year (1-9 divisions per year, **Fig. 2d**). This simple model could explain the differences in SBS1 observed between these two cell types, but does not explain the relative difference in the SBS1 and SBS5/40 mutation rate in seminiferous tubules.

We also explored the existence of DNA damage or repair mechanisms that may vary between normal cell types irrespective of cell division rates and which may be specific to the seminiferous tubules. First, we examined mutation rates in different sectors of the genome (**Methods**), observing a reduced rate in exons compared to introns in most somatic tissues (as previously shown in cancer genomes^41^). This was not, however, found in seminiferous tubules, in *de novo* mutations (DNM) obtained from 100k healthy trios^23^ or in 107 million singleton population variants from GnomAD^42,43^ (**Fig. 4b**). Second, we examined the effect of gene expression level on mutation rate. There was a consistent trend for decreasing mutation burden as expression levels increase in somatic tissues (**Fig. 4c**), with the exception of oesophagus, in which a substantial SBS16 burden in one individual drives increased mutagenesis in highly expressed genes and obscures the effect (**Methods**; **Extended Data Fig. 8**). Seminiferous tubules, however, showed no evidence of a reduction in mutation rate with increasing gene expression, a similar pattern to DNMs and GnomAD^33^. Furthermore, our data suggest that the ratio of SBS on the transcribed and non-transcribed strands in the seminiferous tubules is similar to that in somatic tissues (**Fig. 4a**). Hence, transcriptional coupled repair (TCR) does not seem to be exclusive to the germline and is similar to other somatic tissues studied here, contrary to a previous proposal^44^. Third, we examined the effects of replication timing on mutation rate and found enrichment of mutations in late replicating regions in all cell types, primarily driven by SBS1 and SBS18 mutations (**Fig. 4d; Extended Data Fig. 9**). However, this was much weaker in seminiferous tubules than in somatic cells. Overall, the results suggest that factors other than cell division rate, relating to the intrinsic biology of germline cells, also contribute to differences in mutation load between somatic cells and the germline.

**Figure 4.**
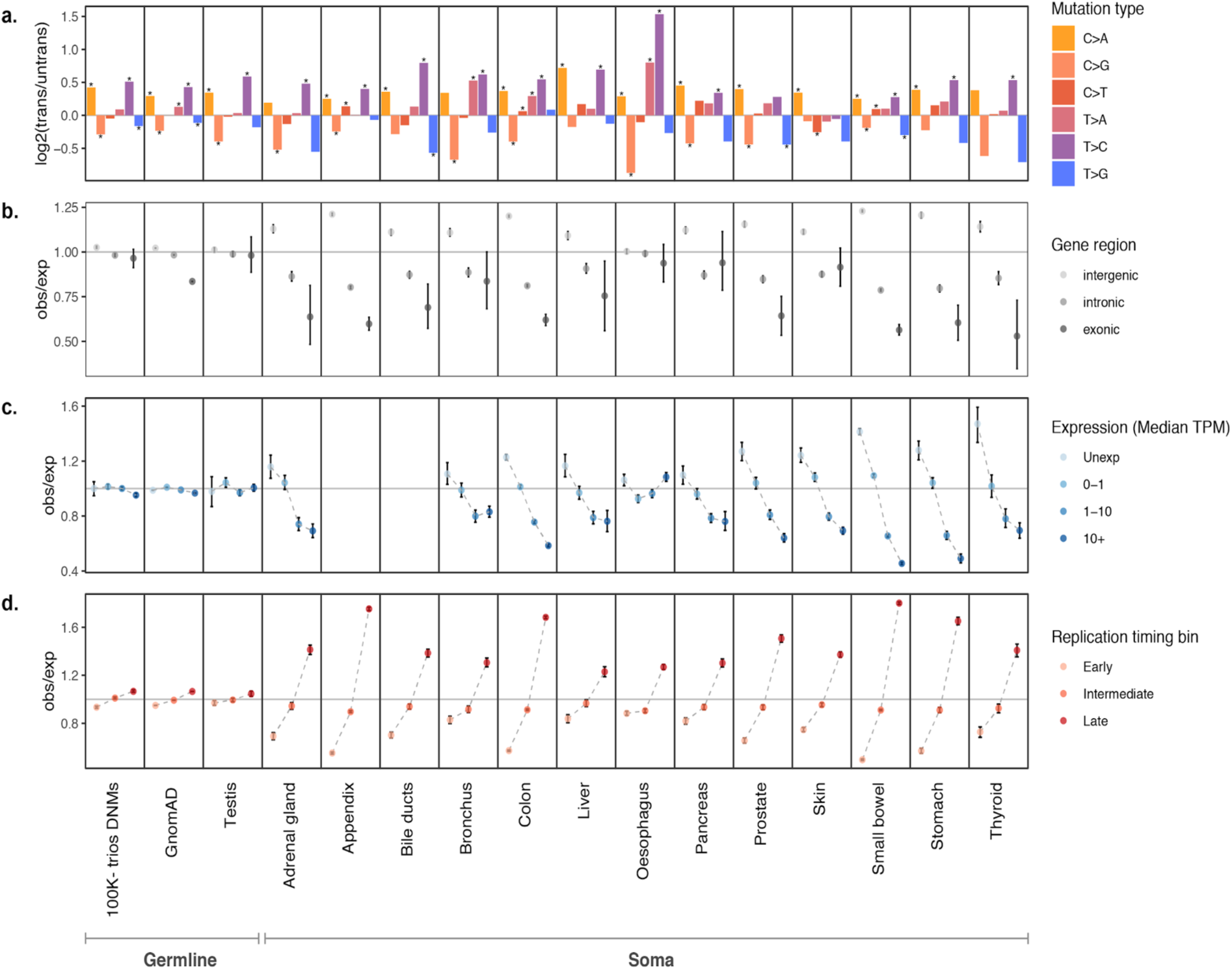
Comparison of mutational biases between the germline and soma. Three SNV germline variant datasets were compared with 13 somatic tissues. **a.** The log2 ratio of SNVs on the transcribed to non-transcribed strands for the 6 mutation classes. Asterisks indicate significant transcriptional strand biases after accounting for multiple tests (P < 0.05, two-sided Poisson test). **b-d.** Observed/expected mutation burden for **b**. Intergenic, intronic, and exonic regions **c.** Tissue-specific gene expression level bins, and **d**. Early, intermediate, and late replicating regions of the genome. The expected burden for a bin is calculated based on the trinucleotide counts of regions in that bin and the average trinucleotide mutation rates in that tissue.

## Discussion

Using multiple samples from the same individuals, we have compared clonal structures, mutation rates and mutational signatures across an extensive range of normal cell types, substantially extending results from previous studies^2,9–11,13,45,46^. Our results revealed the extent of variation in clonal dynamics across tissues. Most tissues are mosaics of clones originating from single stem cells and many microscopically visible glandular structures are derived from recent single stem cell ancestors. Mutation rates vary between different cell types, with stem cells of the small and large intestinal epithelium exhibiting the highest mutation rates thus far reported (except for skin). Several mutational signatures were observed among normal cell types. However, most mutations in almost all cell types were due to the ubiquitous signatures, SBS1 and SBS5/40. Both are thought to be due to endogenous mutagenic processes. The relative contributions of these signatures differed between normal cell types, indicating that their rates of generation are, at least partially, independently regulated.

Patches of cells dissected from seminiferous tubules of the testis were frequently monoclonal, indicating that they arise from single spermatogonial stem cells. The mutation burdens of these germ cell clones were ~27 fold lower than colorectal crypts from the same individuals and their mutation burdens accumulated with age in a linear manner at 2.02 mutations/year (CI95% 1.83-2.42), lower than all other somatic cells thus far estimated^2,9–11,13,45,46^. How the male germline achieves this low mutation rate has remained elusive. Spermatogonial stem cells face a dilemma; they are under evolutionary pressure to minimise potentially deleterious mutations, yet they must constantly proliferate to maintain spermatogenesis, a process thought to be inherently mutagenic. The results therefore quantify the extent to which the germline is protected from mutations compared to the ‘disposable soma^47^. The haploid mutation rate in seminiferous tubules was strikingly consistent with the paternal contribution to *de novo* germline mutations and the age effect inferred from trio studies^17,19,23^. The results indicate that the low germline mutation rate is not the result of a genetic bottleneck or of selection against mutations during conception or development but is an intrinsic feature of the male germline compared to the soma.

The mutational signatures present in seminiferous tubules were also present in somatic cells, predominantly SBS1 and SBS5/40, but with lower mutation rates of both signatures compared to other somatic cells. The lower burden in the seminiferous tubules was predominantly due to the lower rate of SBS5/40, which accounts for most mutations in both cell types, but with a much greater reduction in SBS1. At least in part, these differences could be due to a lower rate of spermatogonial stem cell division compared to somatic stem cells. However, differences between seminiferous tubules and somatic cells in the distribution of mutations across the genome suggest that other intrinsic biological differences may also play a role^39^. It is thus possible that superior mechanisms of DNA maintenance may cause a lower mutation rate per cell division and thereby contribute to the lower mutational burden observed in the germline compared to the soma.

This first survey of mutations acquired during lifetime in multiple somatic and germline cells from the same donors advances our understanding of the diversity of mutation rates and processes within the human body. We quantify a uniquely low mutational burden in the germline relative to somatic cells. We find that the mechanisms underlying mutagenesis appear to be shared between the germline and the soma, suggesting that the germline has found ways to limit the mutagenesis caused by these processes. While our analyses hint at what some of these mechanisms may be, further work will be necessary to elucidate precisely how the germline protects itself from the considerably higher mutation rates observed in the soma.

## Methods

### Sample collection

Anonymized snap-frozen samples were retrieved from living and deceased donors. These are outlined below.

### Panbody tissue samples

#### Donor PD28690

Multiple samples from 22 macroscopically normal tissues and organs (**Supplementary Table 1**) were collected from a 78-year-old male during a rapid autopsy (rapid autopsy defined as an autopsy with a post-mortem time interval (PMI) of < six hours). This donor was a non-smoker who died of a metastatic oesophageal adenocarcinoma for which he had received a short course of palliative chemotherapy (5-6 weeks of oxaliplatin 7 weeks prior to death). He had no other comorbidities. The samples were collected in line with the protocols approved by the NRES Committee East of England (NHS National Research Ethics Service reference 13/EE/0043). Every sampled tissue was photographed and biopsy sites carefully documented. Once collected, all tissue biopsies were snap frozen in liquid nitrogen and subsequently stored at −80°C. Summary of all sampled tissues is provided in **Supplementary Table 2**.

#### Donors PD43850 and PD43851

Multiple biopsies from 16different tissues (**Supplementary Table 1**) were collected from a 54-year-old female (PD43850) and a 47-year-old male (PD43851/PD42565); both individuals died of non-cancer causes (traumatic injuries and acute coronary syndrome respectively). Similarly, all samples were obtained within less than six hours of death (one hour and three hours respectively), were snap frozen in liquid nitrogen and subsequently stored at −80°C. The use of these tissues was approved by the London, Surrey Research Ethics Committee (REC reference 17/LO/1801, 26/10/2017).

#### Additional testis and colon samples

AmsBio (commercial supplier) – Samples for the following donors: PD40744, PD40745/PD42564, PD40746/PD42568, PD42563, PD42566, PD42569 were obtained at autopsy from individuals who died on non-cancer related causes (**Supplementary Table 1**). The use of these tissues was approved by the London, Surrey Research Ethics Committee (REC reference 17/LO/1801, 26/10/2017). Mr. Rakesh Heer (consultant surgeon) - Samples for the following donors: PD42036, PD42034, PD43727, PD43726, PD46269 were obtained from individuals who had non-testicular problems such as abdominal chronic pain and tissue distortion (**Supplementary Table 1**). The use of these samples was approved by North East Newcastle & North Tyneside1 (REC reference 12/NE/0395, 22/09/2009). As above, all collected samples were snap frozen in liquid nitrogen and subsequently stored at −80°C.

#### Laser capture microdissection of tissues

We aimed to explore somatic mutations in relatively small populations of cells from specific morphological or functional units, such as endometrial glands or colonic crypts (**Fig. 1a**). These units typically contain 200-2000 cells. All tissue biopsies were received fresh frozen. A mixture of frozen and paraffin embedded tissue sections were used for laser-capture microdissection (LCM). For frozen preparations, tissues were embedded in an optimal cutting temperature (OCT) compound. 14 to 20-micron thick sections were generated at −20°C to −23°C, mounted onto poly-ethylene naphthalate (PEN)-membrane slides (Leica), fixed with 70% ethanol, washed twice with phosphate-buffered saline (PBS), and stained with Gill’s haematoxylin and eosin for 20 and 10 seconds respectively.

For paraffin preparations, the above outlined frozen tissues were first thawed at 4°C for 10-15 minutes. They were then fixed either in 70% ethanol or Paxgene (PreAnalytiX, Hombrechtikon, Switzerland) and embedded in paraffin using standard histological tissue processing (Ellis et al, 2020). 8 to 10-micron thick sections were subsequently cut, mounted on to PEN-membrane slides, and stained by sequential immersion in the following: xylene (two minutes, twice), ethanol (100%, 1 minute, twice), deionised water (1 minute, once), Gill’s haematoxylin (10-20 seconds), tap water (20 seconds, twice), eosin (10 seconds, once), tap water (10-20 seconds, once), ethanol (70%, 20 seconds, twice) and xylene or neoclear xylene substitute (10-20 seconds, twice).

Using the LCM (Leica LMD7), each biopsy was first examined and the benign nature of the tissue was confirmed. Specific cell populations or microscopic structures were then visualised, dissected and collected into separate wells in a 96-well plate. Overview, pre- and post-dissection images were taken. Haematoxylin and eosin (H&E) histology images were subsequently reviewed by two pathologists (LM, TRWO, MJL, AYW). Tissue lysis was performed using Arcturus PicoPure Kit (Applied Biosystems) as previously described^1,2^, Ellis et al, 2020.

#### Microdissection of human tissues

Each patch included part or all of individual colorectal, appendiceal and small intestinal crypts; prostatic and gastric glands; pancreatic acini, ducts and islets; thyroid follicles; biliary, renal and seminiferous ducts/tubules; segments of full thickness squamous epithelium from the skin and oesophagus; urothelium from the bladder and ureter; sero-mucous glands and pseudostratified epithelium from bronchus; and patches without microanatomical structure from within liver parenchyma, cardiac muscle, adrenal cortex, adipose tissue and brain. (**Supplementary Table 2**)

#### Library preparation and whole genome sequencing

The lysate was submitted to a bespoke low-input library pipeline without prior DNA quantification, as has been previously described^1^. Briefly, the method utilises enzymatic fragmentation and PCR amplification to generate sufficient libraries from single cuts of colonic crypts and seminiferous tubules for whole genome sequencing. These libraries were then used to generate 150bp paired end sequencing reads on either Illumina XTEN or NovaSeq platforms. The target coverage was ~30x.

#### Variant calling

Sequencing data were aligned to the reference human genome (NCBI build 37) using Burrows-Wheeler Aligner (BWA-MEM)^3^. Duplicates were marked using biobambam2^4^.

#### Substitutions

Single base somatic substitutions were called using the CaVEMan algorithm (major copy number 5, minor copy number 2)^5^. To exclude germline variants for investigation of cellular mechanisms of low mutation burden, matched analysis was performed using bulk sequencing of matched tissue (blood, skin, fat or colon).

Further post-processing filters were then applied. Common single nucleotide polymorphisms (SNPs) were first filtered against a panel of 75 unmatched normal samples before recurrent artefactual mutations associated with the aforementioned low-input DNA sequencing pipeline were eliminated using previously validated fragment quality thresholds. Additional filters were applied to remove mapping artefacts associated with BWA-MEM. The median alignment score of reads that support a mutation should be greater than or equal to 140 (ASMD ≥ 140) and fewer than half of the reads should be clipped (CLPM = 0).

The resultant variant list provided a list of sites for genotyping across each patient to maximise mutation detection. A count of mutant and wildtype bases at each site was generated with mapping quality threshold of 30 and base quality threshold of 25 necessary to count a base as mutant. Using only samples from the derived count matrix confirmed to be diploid following copy number review (described below), we applied an exact binomial test to filter residual inherited variants and a maximum-likelihood estimation of the beta-binomial overdispersion to flag and remove remaining mapping artefacts^6^. The cut-off for the overdispersion parameter (rho) was set to 0.1, as done previously^7^. Substitutions common across multiple (>2) samples from the same individual were manually reviewed and low-quality calls were removed.

#### Indels

Insertions and deletions were called using the cgpPindel algorithm followed by standardised in-house post processing^8^. As before, samples were run against matched-normals to avoid calling germline variants. LCM samples exhibited high numbers of artefactual insertion variants occurring at homopolymer runs of 9+ and thus these were filtered out. In addition, based on allele counts estimated by exonerate^9^, variants with ≥20% VAF in the matched normal sample and variants with VAF in matched normal samples greater than the VAF in the sample were filtered out. Variants overlapping genomic locations with mean coverage < 10 and > 100 across samples were further filtered out. Similar to the approach detailed for substitutions, the candidate sites were then genotyped across all patient samples and underwent the exact binomial and beta binomial filtering (rho cut-off 0.2) to exclude common variants and systematic artefacts calls.

Based on the manual review of the remaining insertions and deletions, additional filters were applied to exclude, a) Common variants called across multiple individuals b) Variants occurring on reads with a greater frequency of ambiguous and unknown genotypes than the reads with the reference allele c) Variants at positions where the ratio of good quality reads (MQ ≥20) to all reads < 0.5.

#### Copy number and structural variants

Copy number variants were called with Allele-Specific Copy number Analysis of Tumours (ASCAT) using matched-normals. Identified CNVs were manually inspected; first by review of the associated logR and B-allele frequency (BAF) plots and then by visual inspection of the raw coverage across that region of the genome.

Filtering called SVs was performed as done previously^2^ via AnnotateBRASS v3 (https://github.com/MathijsSanders/AnnotateBRASS). In brief, for each sample an appropriate set of control samples is defined for filtering SVs on a subset of calculated metrics. A control sample is considered appropriate when phylogenetically unrelated to the sample of interest and lacking any recurrent SVs. The batch primarily comprised polyclonal samples (e.g., brain or stroma) without evidence of detectable clonal composition. Metrics were calculated as described previously^2^ and the same filtering approach was executed. SVs indicative of translocations were reviewed and reclassified as part of retrotransposon (RT) insertions when strong hallmarks of the latter was present. These included proximal breakpoints on both chromosomes indicative of a small sequence insertion, known RT source hotspots, multiple events stemming from a RT source hotspot in the same sample and the presence of long polyA/T-tails at inserted sites.

Coverage and BAF information were extracted for all samples by ConstructASCATFiles (https://github.com/MathijsSanders/ConstructASCATFiles). Single nucleotide polymorphisms (SNPs) with a minor allele population frequency greater than 0.01 were used as positions for extracting the aforementioned information. Coverage and BAF information were grouped by donor and assessed for quality via the ‘QualityControl_and_PCA.R’ script (https://github.com/MathijsSanders/PREASCAT). In brief, for each donor one or more control samples were designated which are assumed to comprise cells possessing a normal karyotype (i.e. normal stromal tissue). SNPs with limited coverage across the control samples are excluded from analysis. Samples are corrected for library size and the LogR ratio is determined by comparing the coverage of each sample to the median coverage across the batch of predefined controls. For male individuals the coverage of chromosome X is multiplied by 2 to correct for sex differences. The BAF profile of the germline is determined by taking the median BAF for each SNP across the batch of control samples belonging to the same individual. This step is to maximize the identification of heterozygous SNPs due to higher BAF noise for low-input protocols. Principal component analysis (PCA) was applied to identify systematic biases present across all samples included. The first principal components (PCs) primarily represent regions of high coverage variability or high levels of polymorphism (e.g. the HLA locus), high-versus-low CpG density and, in rare cases, regions of open chromatin. This information is exported into WIG or bigWig format for review in a genome browser. Formal LogR and BAF values were calculated via the ‘construct_ASCAT_files.R’ script (https://github.com/MathijsSanders/ConstructASCATFiles). The procedure is similar to the above with the exception that the PCs are used in a linear regression setting. Coverage profiles are centred around mean 0 and the PCs are regressed against the 0-centered coverage profile. The fit is subtracted from the 0-centered coverage profile and the profile is restored to its original mean values. This procedure removes most of the biases present in the first PCs. Finally, the necessary files are exported for ASCAT analysis. The GC content in monotonically increasing windows centred on the SNPs used in this analysis are calculated by ContentGC (https://github.com/MathijsSanders/ContentGC). Generated input files and the GC-content file were used in ASCAT per default.

#### Observed vs. Expected Mutation Burden

To calculate expected mutation burden for a tissue type in a region of interest (e.g. replication timing bin, set of genes) we first computed the overall mutation rate of the 32 possible trinucleotide contexts within that tissue. To determine the mutation rate for each context, we divided the occurrences of SNVs at that trinucleotide by the total observations of that trinucleotide within the callable genome of the tissue. The expected number of variants for a given trinucleotide context is the mutation rate of that context multiplied by the total observations within the region of interest. The expected mutation burden is given by summing the expected variants from each of the 32 trinucleotide contexts.

Confidence intervals for observed/expected mutation burden were generated by bootstrapping with 10,000 random samplings with replacement of observed mutations and recalculations of observed to expected ratios (https://github.com/Rashesh7/PanBody_manuscript_analyses/Signature_Enrichment). Expression levels were tissue specific median transcripts per million (TPM) from Genotype-Tissue Expression (GTEx) project v8 data^10^. Replication timing values were median values of 1-kb genomic bins across16 ENCODE project cell lines^11^ that were divided into early (≥60), intermediate (>33 & <60) and late (≤33) bins as previously described.

#### Clonal decomposition

The clonality of LCM samples was assessed through a truncated binomial mixture model. Truncated binomial distributions were used as a basis rather than regular binomial distributions to account for the censoring of variants below four supporting reads, which is hard-coded in CaVEMan. In effect, the binomial distributions are renormalised to the observable distribution to arrive at their truncated counterparts. Using expectation-maximisation, between one and five clones were fitted to the distributions of number of variants supporting reads and total depth per LCM sample. The optimal decomposition was chosen through the Bayesian information criterion (BIC). The mixture model then yields the underlying probability (VAF) and proportion for each component. A component was defined to be clonal if the VAF was higher than 0.25, i.e. the clone pervades the majority of cells in the sample. This ensures that the clone is composed of a single lineage, since no two independent clones can overlap at VAF>0.25, which would account for more than the total of cells. The estimated proportion of variants attributable to clonal components was then used to correct the observed mutation burden.

#### Mutational signature analysis

Mutational signatures were extracted using two algorithms 1) HDP (https://github.com/nicolaroberts/hdp) based on the Bayesian hierarchical Dirichlet process 2) SigProfiler (https://github.com/AlexandrovLab) based on non-negative matrix factorisation.

HDP was run without priors on single base substitutions (SBS) derived from phylogenetic branches rather than samples to avoid double-counting shared mutations (ref to phylogeny paper). Branches with fewer than 100 total mutations were excluded from the extraction.

In this way, fourteen signature components were extracted (Supplementary Information). A subset of these fourteen signatures appeared to be combinations of previously reported reference signatures. To deconvolute composite signatures and to equate obtained HDP signatures to reference signatures, we employed an expectation maximisation-algorithm to deconstruct these signatures into reference constituents. The set of reference signatures included was informed by an HDP run with references signatures from COSMIC v3.1 as priors. If any priors remained in the final call set for HDP in this run, they were included in the set of candidate signatures for deconvolution of *de novo* signatures. This amounted to reference signatures SBS1, SBS2, SBS4, SBS5, SBS7a, SBS7b, SBS13, SBS16, SBS17b, SBS18, SBS22, SBS23, SBS32, SBS35, SBS40, SBS41 and SBS88.

In this way, the extracted signatures were deconvoluted into the following reference signatures.

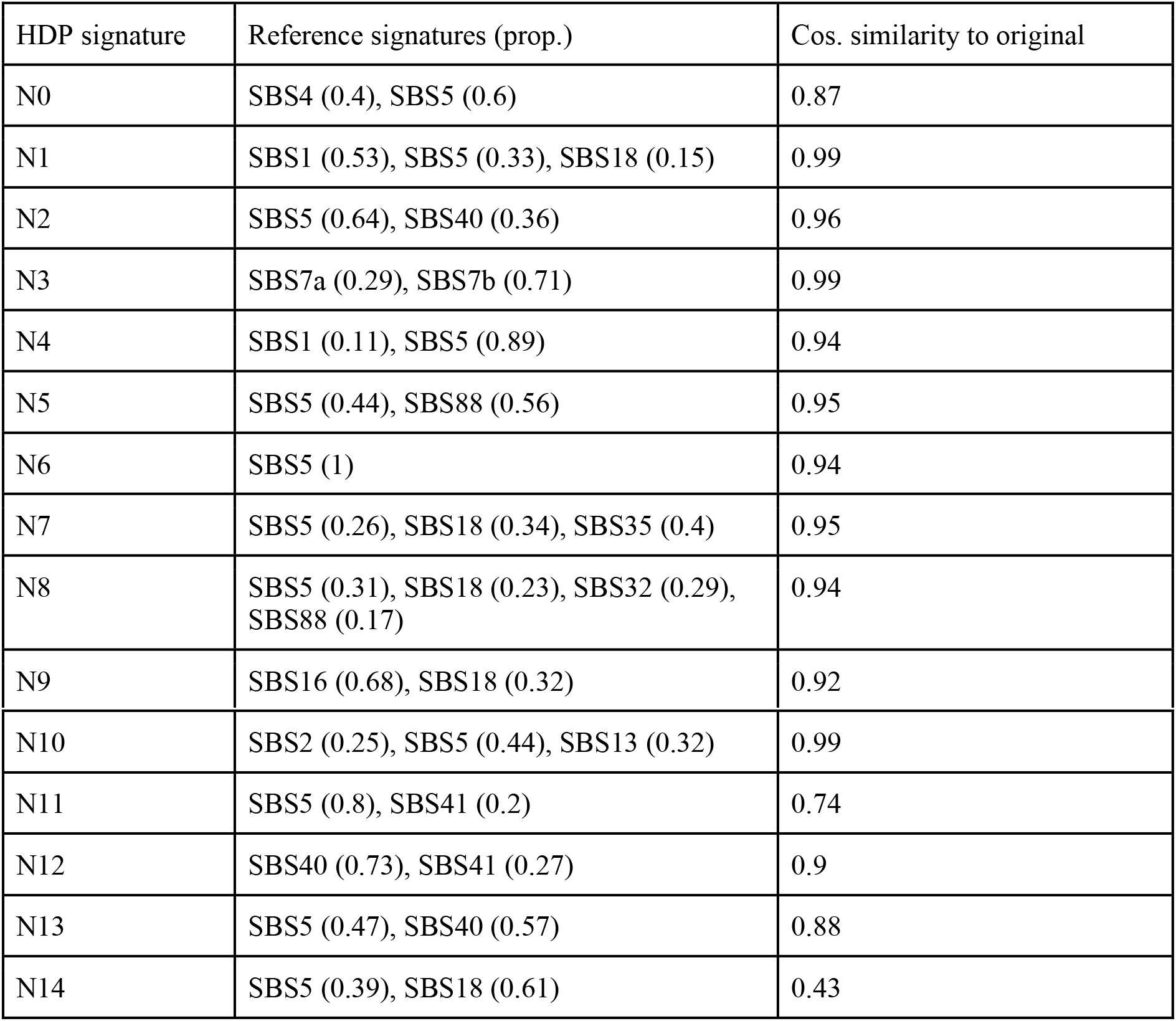

Because of the low cosine similarity (<0.85) with the reconstructed signatures after deconvolution, N11 and N14 were taken forward as a *de novo* mutational signature. Signatures were then fitted to mutational count data per sample. The signatures to be fitted to individual tissues were selected based on the results of the HDP run, to avoid overfitting. If an HDP signature was responsible for more than 7.5% of substitutions in more than two samples, its deconvoluted reference signatures were included in the set of candidate signatures for that tissue. The signature fitting was performed using the SigFit algorithm.

Signature extraction was performed using SigprofilerExtractor V1.0.15, which internally used SigprofilerMatrixGenerator V1.1.18 for generating trinucleotide count matrix^12^, generating solutions between one and 20 de-novo signatures. A solution with 4 de-novo signatures was chosen as the optimal solution, which was further deconvoluted into known COSMIC v3.1 signatures. Other stable solutions with five and six de-novo signatures were also deconvoluted to test the specificity and sensitivity of the solutions, with the four signature solution being more specific and the six signature solution being more sensitive (https://github.com/Rashesh7/PanBody_manuscript_analyses/SigProfiler_analyses). HDP was found to be more sensitive to signatures with smaller contributions and was selected as the main signature extraction method. SBS5 and SBS40 are difficult to deconvolute separately and are usually contaminated by each other during signature extraction. Hence, they were collated together. MutationalPatterns was used to generate the Indel signatures (https://github.com/UMCUGenetics/MutationalPatterns)^13^. Due to low number of insertions and deletions only signatures with indel main types were generated.

#### Mutational burden and age correlation

Samples with median VAF ≤ 0.3 were excluded from burden analyses. Samples were run through a truncated binomial algorithm to identify and allocate mutations to the major clone (VAF ≥ 0.25) as described above^14^. All mutation burden analyses were done using only the major clone counts for each sample. The counts were further corrected for the callable genome of each sample available to CaVEMan for calling substitutions. For calculating callable genome size for each sample; centromeric, telomeric and known simple repeat regions were subtracted from the genome and only the standard contigs (chromosome 1:22, X, Y) were kept; additionally, we applied a coverage filter of ≥ 4X, based on minimum reads required for variant calling. Correlation between mutation burden with age in testis and colon samples was modelled using the following mixed linear model (https://github.com/Rashesh7/PanBody_manuscript_analyses/burden_analyses):

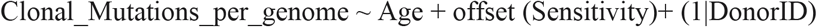

where Sensitivity was calculated as a product of the sample median VAF and sample average coverage.

#### Comparison with Trio based DNMs

The *de novo* mutation counts for 1548 individuals were obtained from deCODE^15^. Paternal *de novo* mutation counts were estimated as 80.4% of the total mutations per sample as estimated in the deCODE^15^ paper. To compare mutation rate per year for testis seminiferous tubules from our study, we estimated haploid SBS counts, by dividing the corrected clonal mutations generated in the above step. The correlation of deCODE^15^ paternal *de novo* mutations with paternal age at conception was modelled by a simple linear model:

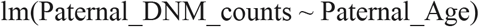

Age correlation of the haploid mutation burden from this study was modeled using a mixed linear model:

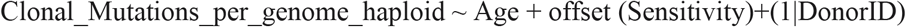

where Sensitivity was calculated as a product of the sample median VAF and sample average coverage. To compare the mutation rate per year between the datasets (paternal *de novo* mutations and haploid clonal burden) a simple linear model was used to query the interaction of the two predictor variables.

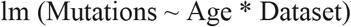

#### Mutation rate per cell division analysis

Assuming that the majority of SBS1 mutations are associated with cell-division we compared the rate of SBS1 per cell division in colonic stem cells and spermatogonia stem cells using the ranges of stem cell divisions per year per tissue. To derive SBS1 mutation rate per tissue per individual, the median of total SBS per tissue per individual was multiplied by proportion of SBS1 (using HDP signature extraction method, above) and divided by the ranges of number of stem cell divisions per year.

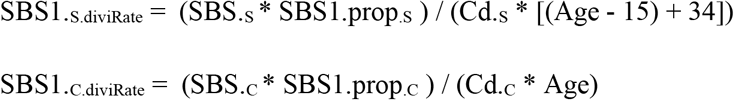

SBS1_.S.diviRate_ is the derived SBS1 mutation rate per cell division in spermatogonial stem cell for a given individual where, SBS_.S_ is the median of SBS in the seminiferous tubules, SBS1_.prop.S_ is the proportion of SBS1 in seminiferous tubules. Age of puberty assumed to be 15 years and the number of cell divisions pre-puberty was estimated to be ~34 (~10 cell division before primordial gemcell (PGC) differentiation + ~24 cell division post PGC). SBS1_.C.diviRate_ is the mutation rate per cell division in colonic crypts stem cells. SBS_.C is_ the median of SBS in colon. SBS1_.prop.C_ is the proportion of SBS1 in colonic crypts. Cd is the number of stem cell divisions per year. A range of spermatogonial stem cell divisions per year was tested for spermatogonia stem cells Cd_.S_ = (365/c (1:10, 16, 20, 25, 30, 35, 40, 45, 50, 60, 80, 100, 125, 180)) and the result was compared to the range of colonic crypts stem cell divisions per year Cd_.C_ = (365/c (60, 80, 100, 125, 180)). Distributions of SBS1 mutation rate per cell division were compared between seminiferous tubules and colonic crypts across ranges of cell divisions per year (above) using the Kolmogorov–Smirnov test. The results showed one to nine divisions per year in spermatogonia stem cells gave a similar rate of SBS1 per cell division to colonic crypts where stem cells divide every two to five days.

#### Telomere length analysis

Telomerecat was used to generate the final telomere length estimates for two main reasons; it provides absolute telomere lengths and is not biased by sequencing depth^10^. The latter point is especially important for our microbiopsies, where sequencing coverage did not uniformly achieve the target 30x depth. For samples which were sequenced using the NovaSeq sequencing platform, the results using Telomerecat were occasionally implausible (such as telomere length estimates of 0bp). NovaSeq sequenced microbiopsies comprised only a small fraction of the cohort (~3%, 19/622), so these samples were omitted for the purposes of telomere analysis.

The Telomerecat values are the average base-pair resolution telomere length for the chromosomes from all cells within a sample. To estimate the effect of age on telomere length in somatic tissues, we used a linear mixed effects model created using the R package lme4 with age and tissue type as fixed effect covariates and patient ID as a random effect. The germline model used the same features, minus the tissue type (all germline sampling was of seminiferous tubules). All models were fitted with an intercept, interpreted as an estimate of telomere length at birth.

#### Detecting Selection

Evidence of selection was assessed using the normalised ratio of nonsynonymous to synonymous substitutions (dN/dS). For a detailed description of this method please refer to Martincorena et al. 2017. The dNdScv (v0.0.1.0) R package^18^ was used to estimate dN/dS. This is a maximum-likelihood implementation of dN/dS that considers mutational biases and is adapted to somatic mutation data. We used the default settings of dNdScv and ran it separately for each cell type, combining all somatic mutations across samples and donors. We also ran dNdScv combining somatic mutations from all samples across all tissues from all donors in order to increase our sensitivity to detect whether any genes were positively selected across tissues. In all cases no genes showed significant evidence of either positive or negative selection after correcting for multiple-hypothesis testing.

#### Identification of Cancer Driver Mutations

To identify any cancer driver variants in these samples we checked whether any of the filtered CaVEMan and Pindel variants occurred within 369 genes previously identified as under selection in human cancers^18^. Variants in these genes were then annotated to indicate mode of action using a catalogue of 764 genes (https://www.cancergenomeinterpreter.org) and a Cancer Gene Census (719 genes)^19^. Truncating variants (nonsense, frameshift and essential splice site) residing in recessive or tumour-suppressor genes were classified as drivers. All missense mutations in recessive or tumour-suppressor genes or in dominant genes or oncogenes were intersected with a database of validated cancer hotspot mutations (http://www.cbioportal.org/mutation_mapper). Missense mutations that occurred in these hotspots were classified as drivers (**Supplementary Table 7**).

## Data Availability

Information on data availability for all samples is available in **Supplementary Table 4**. sequencing data has been deposited in EGA under accession number EGAS00001003021. Substitution, indel, SVs are available in **Supplementary Table 3-6**.

## Code Availability

Pipelines to call SBS, indels, SVs, CNVs, mutation burden analysis, signature extraction with HDP and SigProfiler, mutational burden for different genomic contexts are available from https://github.com/Rashesh7/PanBody_manuscript_analyses.

## Acknowledgements

The authors would like to thank the staff of WTSI Sample Logistics, Genotyping, Pulldown, Sequencing and Informatics facilities for their contribution. We are grateful to: Kirsty Roberts and the cgp-lab for their assistance. We thank Philip Robinson, Martin Goddard, Patrick S. Tarpey, Paul Scott for their assistance with sample collection and LCM-pipeline. We are grateful to Matthew Hurles, Aylwyn Scally and Young Seok Ju for providing useful feedback.

## Funding

This research is supported by core funding from Wellcome Trust. R.R. is funded by Cancer Research UK (C66259/A27114). L.M. is a recipient of a CRUK Clinical PhD fellowship (C20/A20917) and the Jean Shank/Pathological Society of Great Britain and Ireland Intermediate Research Fellowship (Grant Reference No 1175). T.J.M. is supported by Cancer Research UK and the Royal College of Surgeons (C63474/A27176). The laboratory of R.C.F. is funded by a Core Programme Grant from the Medical Research Council (RG84369). Funding for sample collection was through the ICGC and was funded by a program grant from Cancer Research UK (RG81771/84119). R.H. is a recipient of a PCF Challenge Research Award (ID #18CHAL11; Heer). I.M. is funded by Cancer Research UK (C57387/A21777) and the Wellcome Trust. P.J.C. is a Wellcome Trust Senior Clinical Fellow.

## Author Contributions

M.R.S., R.R and L.M. conceived the project. R.R., and M.R.S. supervised the project. A.C., L.M., M.R.S., and R.R. wrote the manuscript; all authors reviewed and edited the manuscript. L.M., A.C., T.C., led the analysis of the data with help from M.D.C.N., R.S., M.S., T.R.W.O., D.L., T.M.B., A.M., K.J.D., R.R. L.M., A.C., and T.R.W.O. performed laser microdissection. L.M. performed the rapid autopsy with help from T.M., and M.T. P.E., Y.H., L.O., and C.L. processed samples. L.M., A.N., R.B., C.I.D, R.H., R.C.F. collected samples. L.M., T.R.W.O., M.J.L., A.Y.W., M.J. reviewed the histological images and clinical reports. I.M., P.C., M.R.S and R.R. helped with data interpretation and statistical analysis.

## Competing Interests

No competing interests are declared by the authors of this study.

---

**Extended Data Figure 1.**
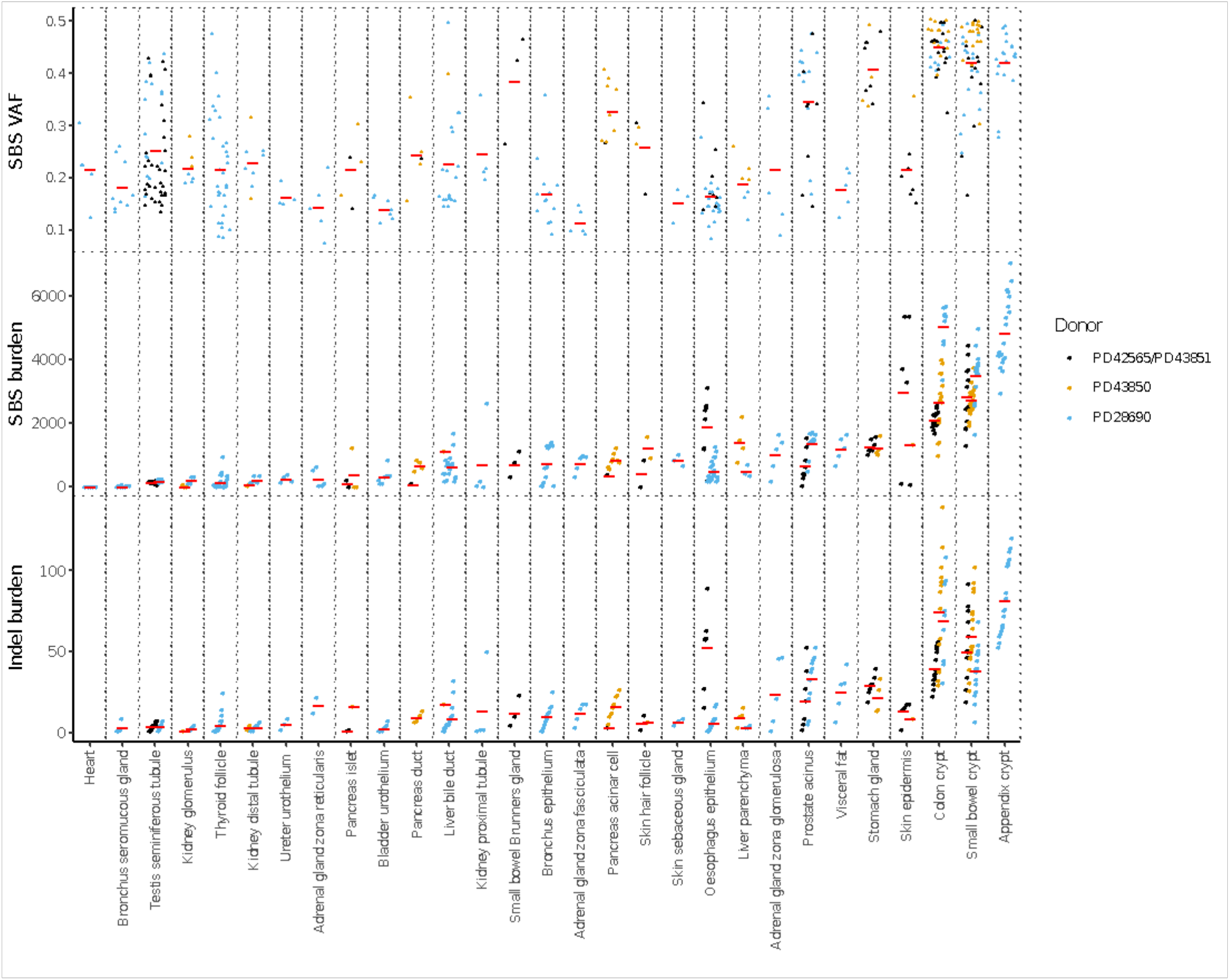
Number of somatic mutations per genome for the 47-year-old male (PD43851), a 54-year-old female (PD43850) and a 78-year male (PD28690) shown by each tissue type. **a**. Median VAF per sample. **b**. SBS burden. **c**. Indel burden.

**Extended Data Figure 2.**
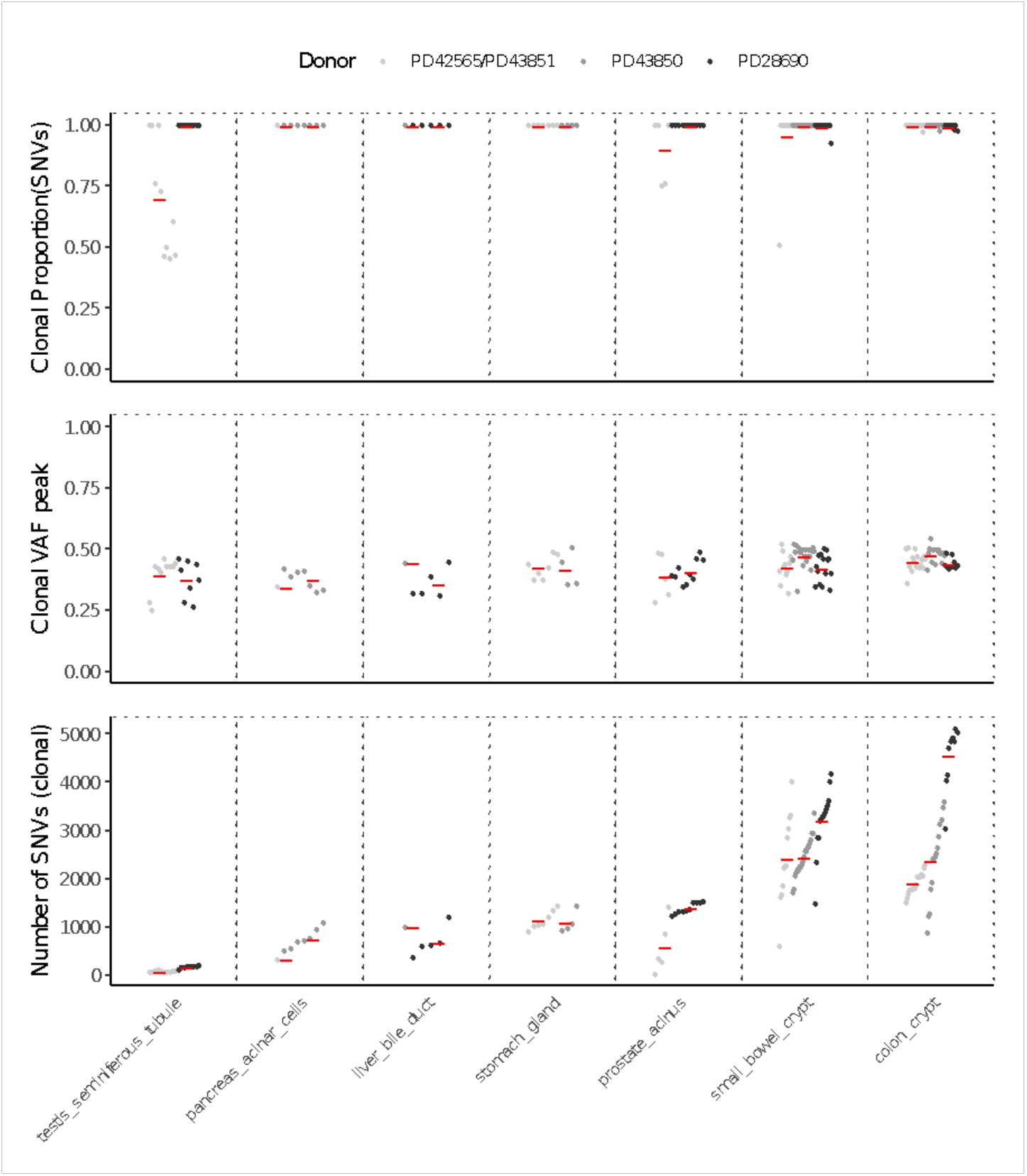
Mutation burden across tissues. Mutation burden was estimated on a subset of tissues that passed all filtering criteria. Minor clone mutations were identified and removed using a truncated binomial algorithm. cell types with a minimum of three samples from more than one individual were included for mutation burden analysis. **a**. Proportion of SBS mutations that were assigned to the major clone by the truncated binomial method. **b**. Peak VAF of SBS belonging to the major clone. **c**. Clonal SBS burden.

**Extended Data Figure 3.**
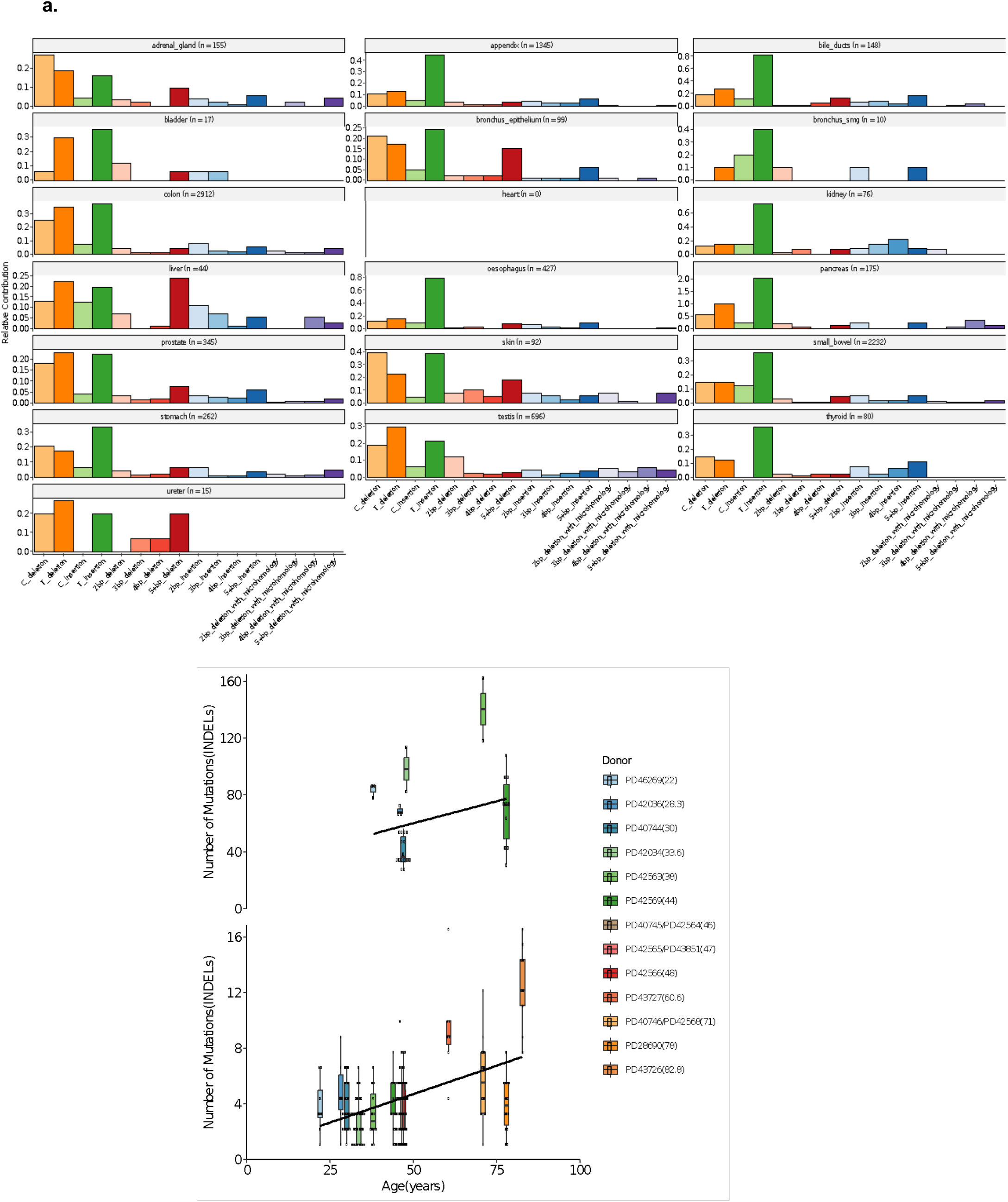
Summary of Indel burden for all 13 individuals. **a**, Indels from each sample were merged together by tissue type. Indel signatures were generated using MutationalPatterns. **b**. Age correlation of Indels per genome (corrected for callable genome) for colon (top panel) and testes (bottom panel). We see a gain of 0.07 Indels per year for testis.

**Extended Data Figure 4.**
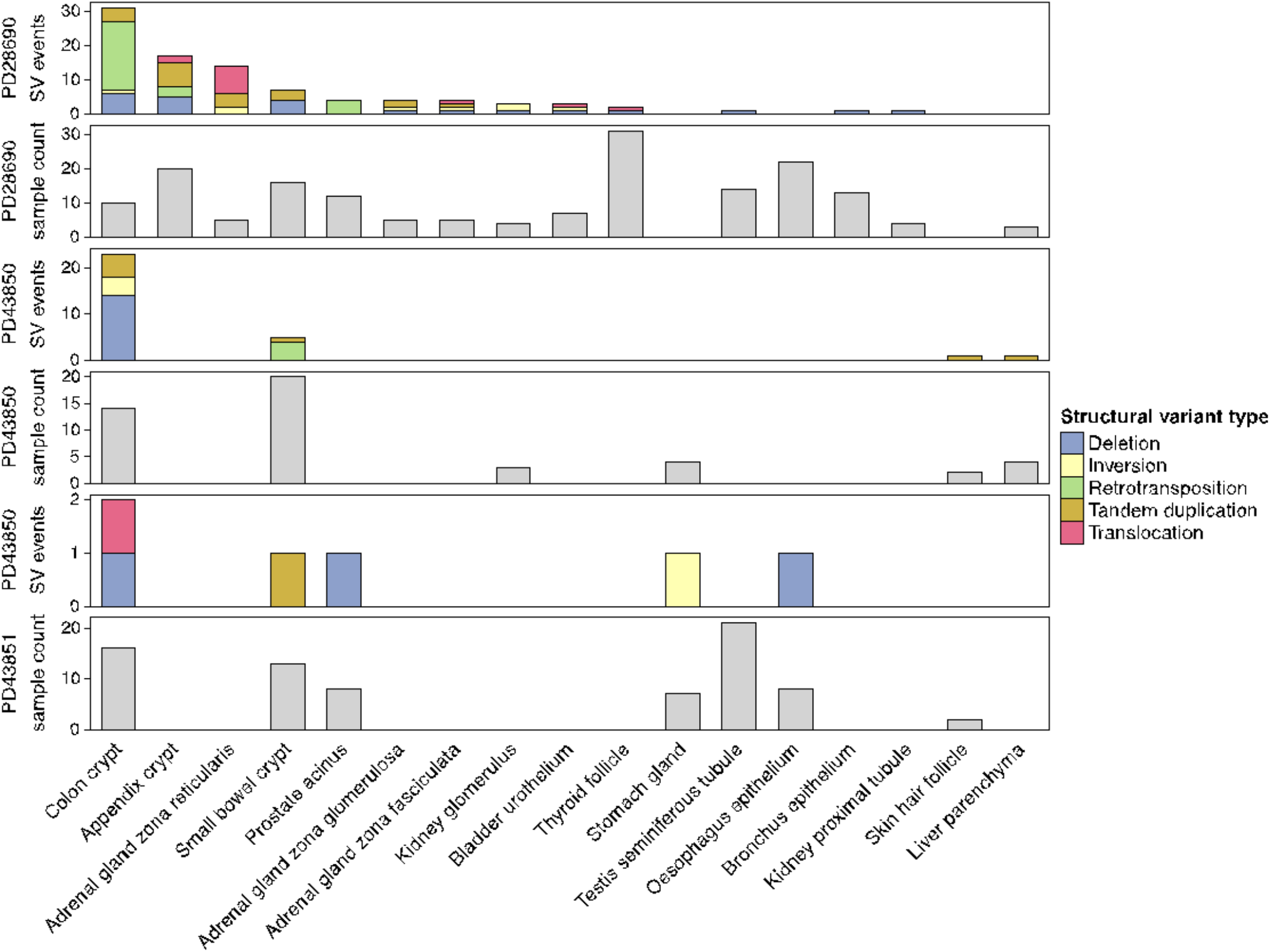
Summary of different structural variation types per tissue per genome. The number of different types of SVs identified, coded by colour, and the number of patches used to identify SV events per tissue per individual. Overall, colonic crypts across all three donors have the highest number of SV events. In particular, high numbers of retrotransposition events were identified in colonic crypts of the 78-year male (PD28690) donor.

**Extended Data Figure 5.**
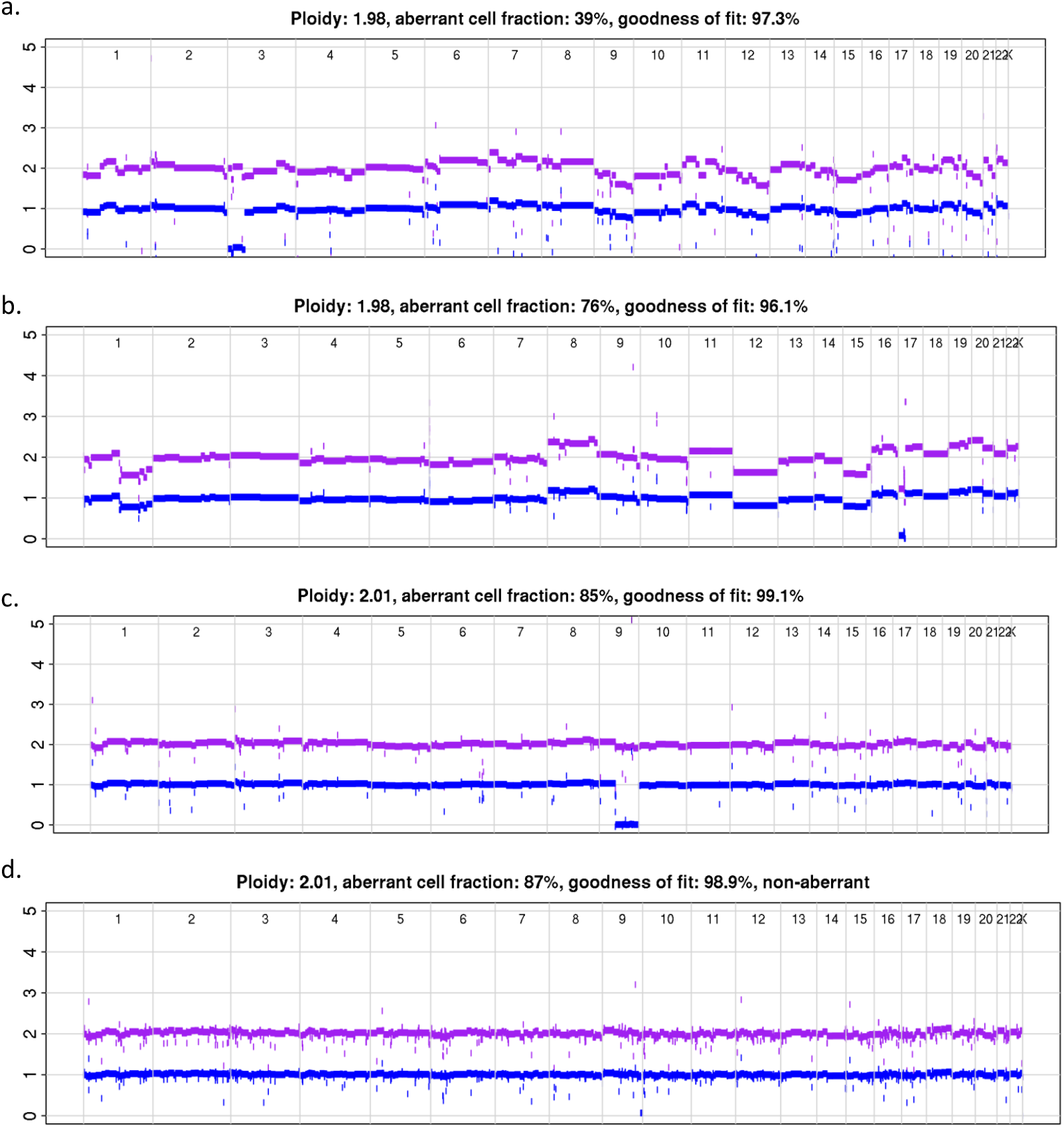
Chromosome-arm or focal losses, encompassing either *NOTCH1* and/or *TP53* in oesophagus. **a.** PD43851k_OSPHG_H12: *NOTCH1* missense mutation (Chr9:Pos139417476:G>T) and subclonal loss 9qter **b.** PD43851k_OSPHG_B2: *TP53* and loss of single copy 17p **c.** PD43851k_OSPHG_E2: *NOTCH1* missense mutation (Chr9:Pos139412332:C>T) and loss 9q **d.** PD43851k_OSPHG_G2: *NOTCH1* missense mutation (Chr9:Pos139412332:C>T) and *TP53* missense mutation (Chr17:Pos7579358:C>A) combined with loss 9qter and *TP53* mutation

**Extended Data Figure 6.**
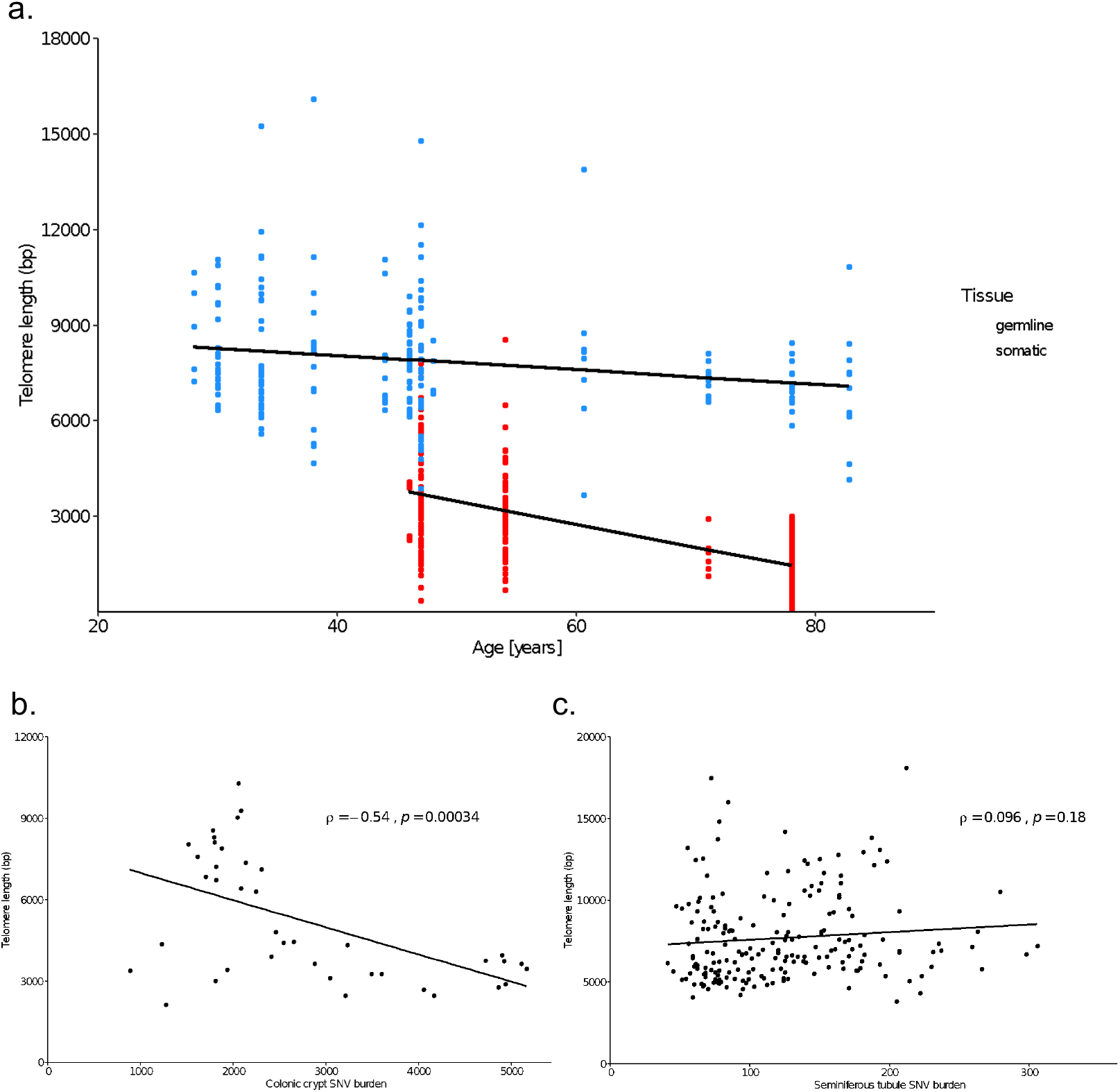
Telomere length comparison between testes and colon. **a.** Regression lines from the linear mixed effects model comparing the impact of age on telomere length between colonic crypts (red) and seminiferous tubules (blue). The points are the partial residuals, controlling for the tissue type. **b.** Correlation between absolute SNV burden and telomere length in the colonic crypt microbiopsies. Spearman’s rank rho and p value stated on plot. **c.** Correlation between absolute SNV burden and telomere length in the seminiferous tubule microbiopsies. Spearman’s rank rho and p value stated on plot.

**Extended Data Figure 7.**
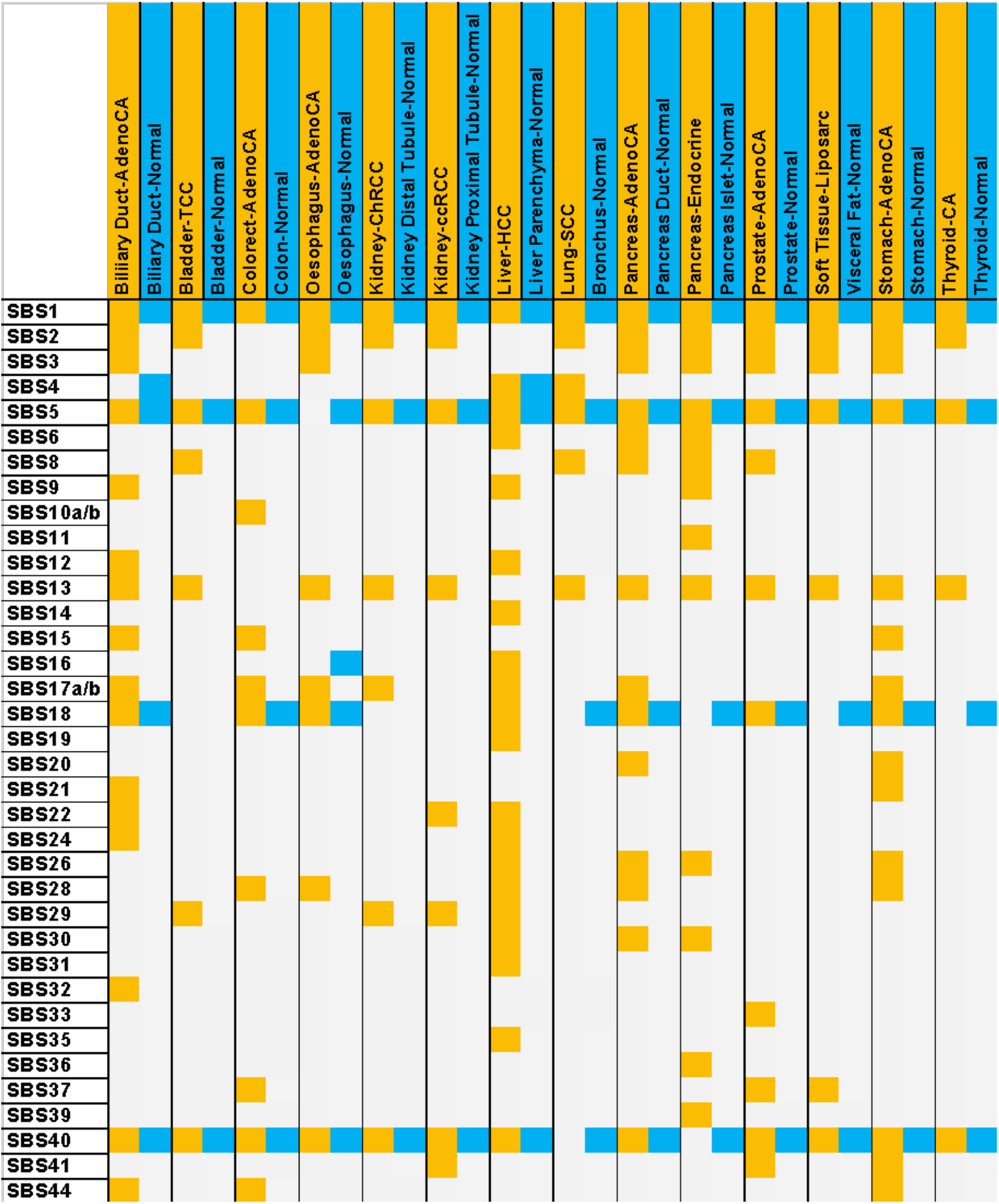
Mutational signatures present in PCAWG tumours compared to normal tissues. The columns contain signatures identified in tumours from the PCAWG dataset adjacent to signatures identified in the corresponding normal tissue type from our dataset^12^. Signatures present in tumours are highlighted in yellow while those present in normal tissue are highlighted in blue.

**Extended Data Figure 8.**
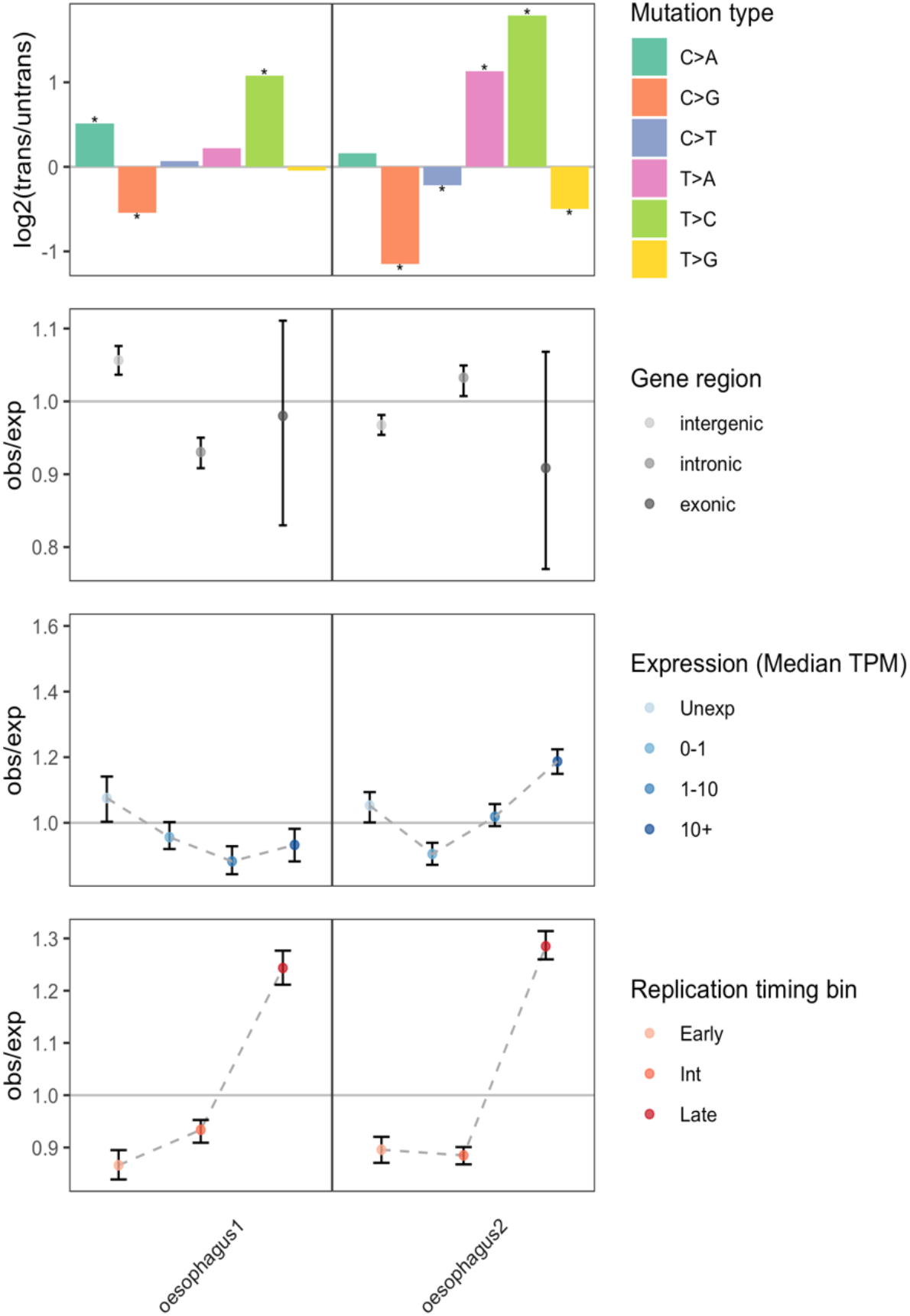
Comparison of oesophagus mutational biases between individuals. a. The log2 ratio of SNVs on the transcribed to non-transcribed strands for the 6 mutation classes. Asterisks indicate significant transcriptional strand biases after accounting for multiple tests (P < 0.05, two-sided Poisson test). b-d. Observed/expected mutation burden for b. Intergenic, intronic, and exonic regions, c. Transcripts across four oesophagus specific GTEx^10^ gene expression level bins, and d. Early, intermediate, and late replicating regions of the genome. The expected burden for a bin is calculated based on the trinucleotide counts of regions in that bin and the average trinucleotide mutation rates in that tissue. Oesophagus 2, PD28690 (78Yrs), with SBS16 shows outlier patterns

**Extended Data Figure 9.**
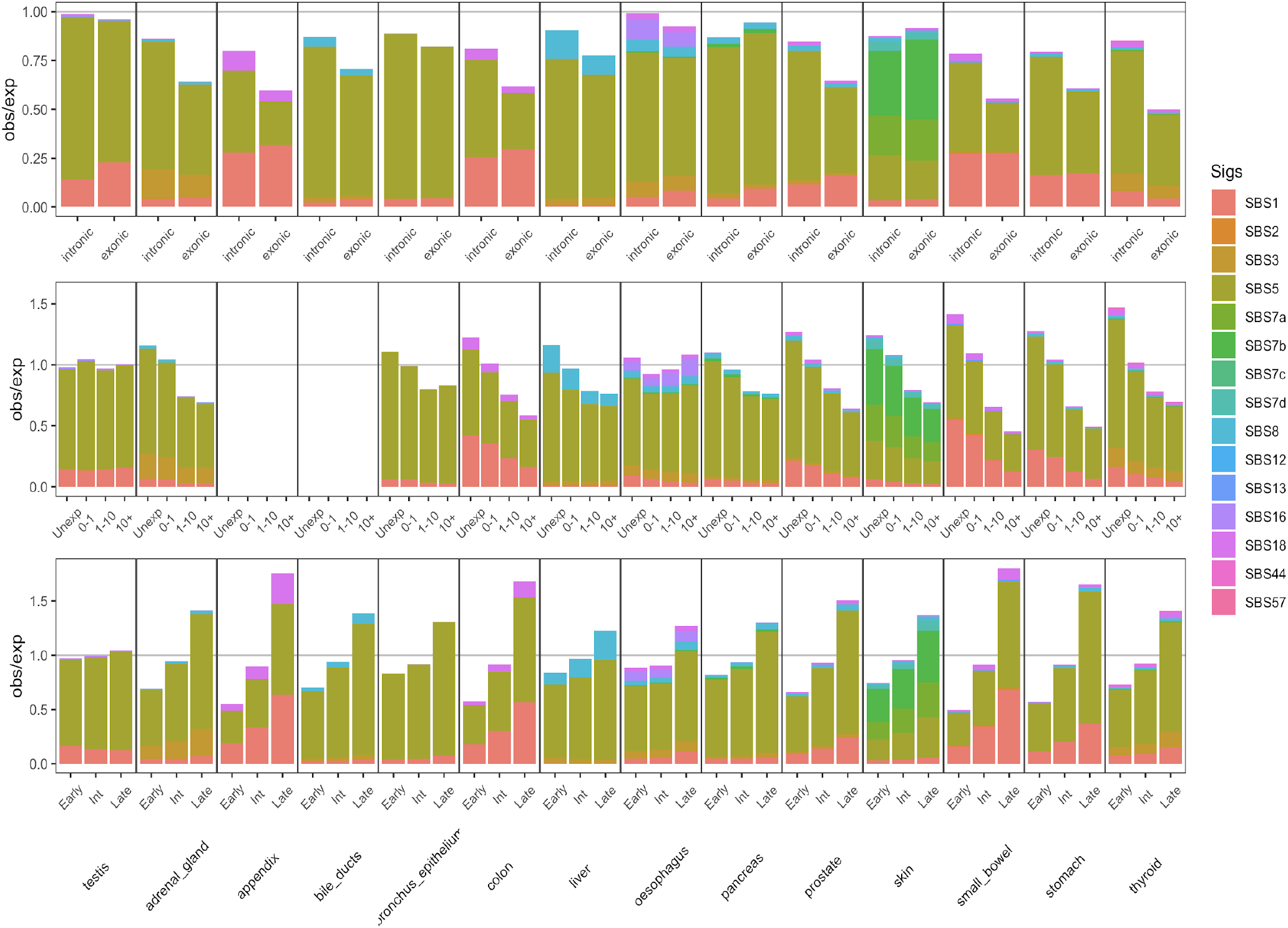
Mutational signature contribution to mutational biases between the germline and soma. **a-c.** Mutational signature contribution to observed/expected mutation burden for a. Intergenic, intronic, and exonic regions, **b.** Transcripts across four tissue specific GTEx^10^ gene expression level bins, and **c.** Early, intermediate, and late replicating regions of the genome. The expected burden for each bin is calculated based on the trinucleotide counts of regions in that bin and the average trinucleotide mutation rates in that tissue. The mutational signature breakdown is calculated using the probability of each variant belonging to each signature based on the fraction of signature in that tissue and the frequency of the mutation type with that signature.

**Supplementary Figure 1.**
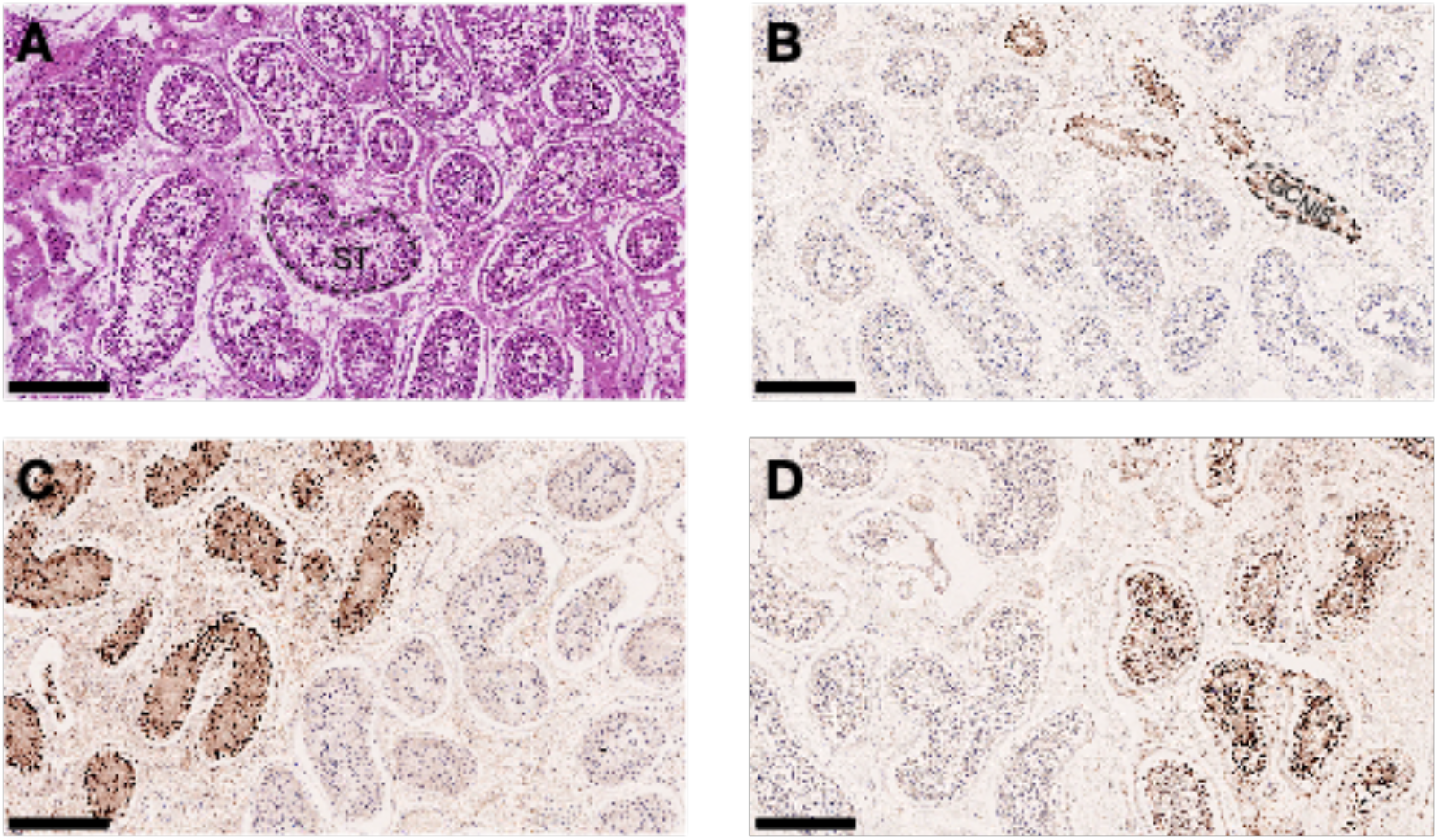
Testicular histology. **a.** H&E stained section from background testicular tissue sampled at the time of tumour resection from PD46269. The majority of the background testicular tissue comprises normal seminiferous tubules (ST) which contain germcells at various stages of maturation. **b-d.** OCT3/4 immunohistochemistry staining of sections of background testicular tissue from the three patients (PD46269, PD42036 and PD42034) that were diagnosed with germ cell tumours. Positive staining for OCT3/4 indicates the presence of germ cell neoplasia situ (GCNIS), the precursor to the invasive tumour. Following review of the H&E stained section morphology and OCT3/4 immunohistochemistry by two histopathologists, we ensured that only microdissected tubules which possessed normal morphology, away from regions of OCT3/4 positivity were included in the analysis. The scale bars denote 250 micrometres.

**Supplementary Figure 2.**
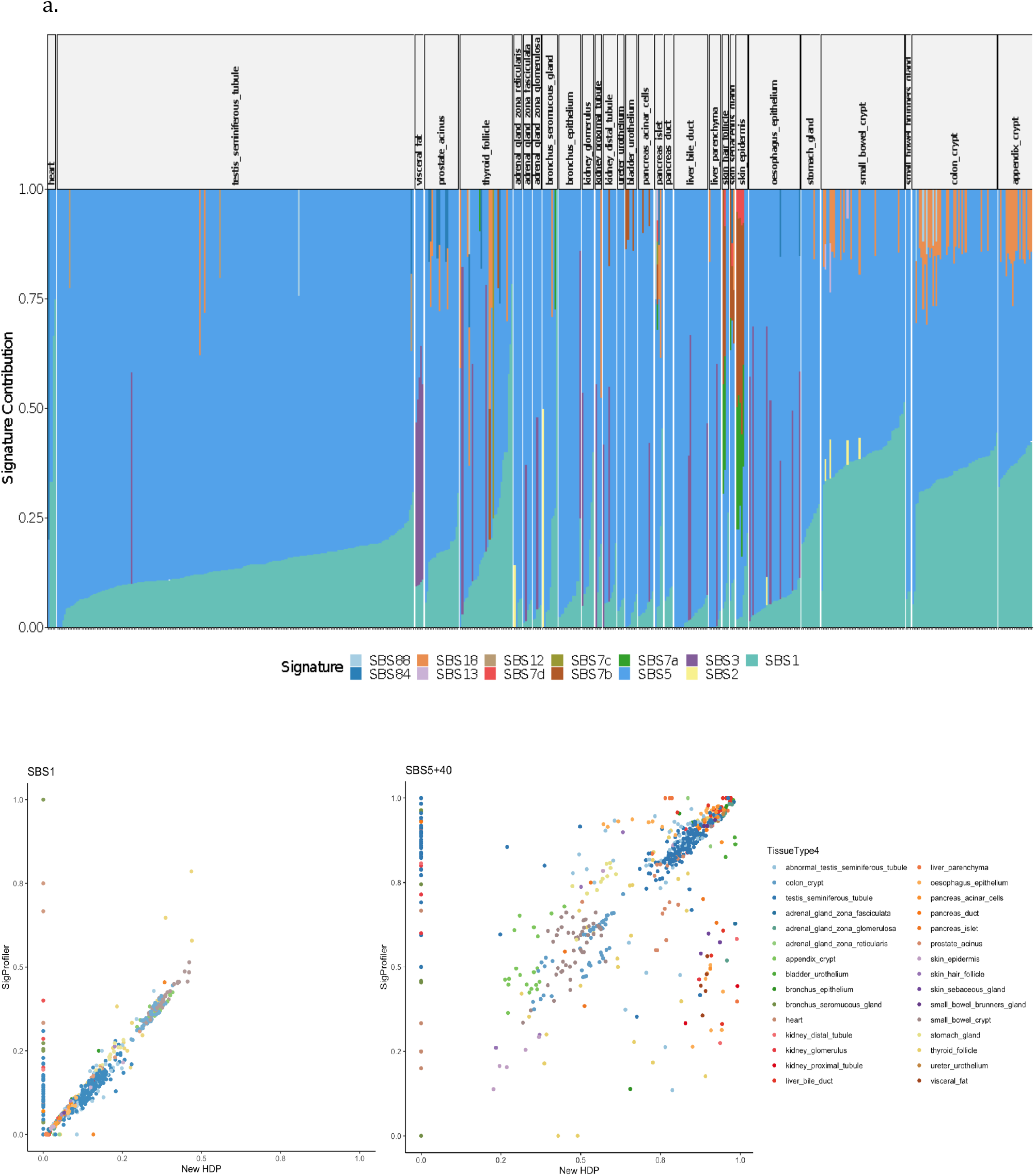
Comparison between Signature extraction methods. **a.** Signature extraction with SigProfiler **b.** Concordance between HDP and SigProfiler in proportion of SBS1 and SBS5/40

**Supplementary Figure 3.**
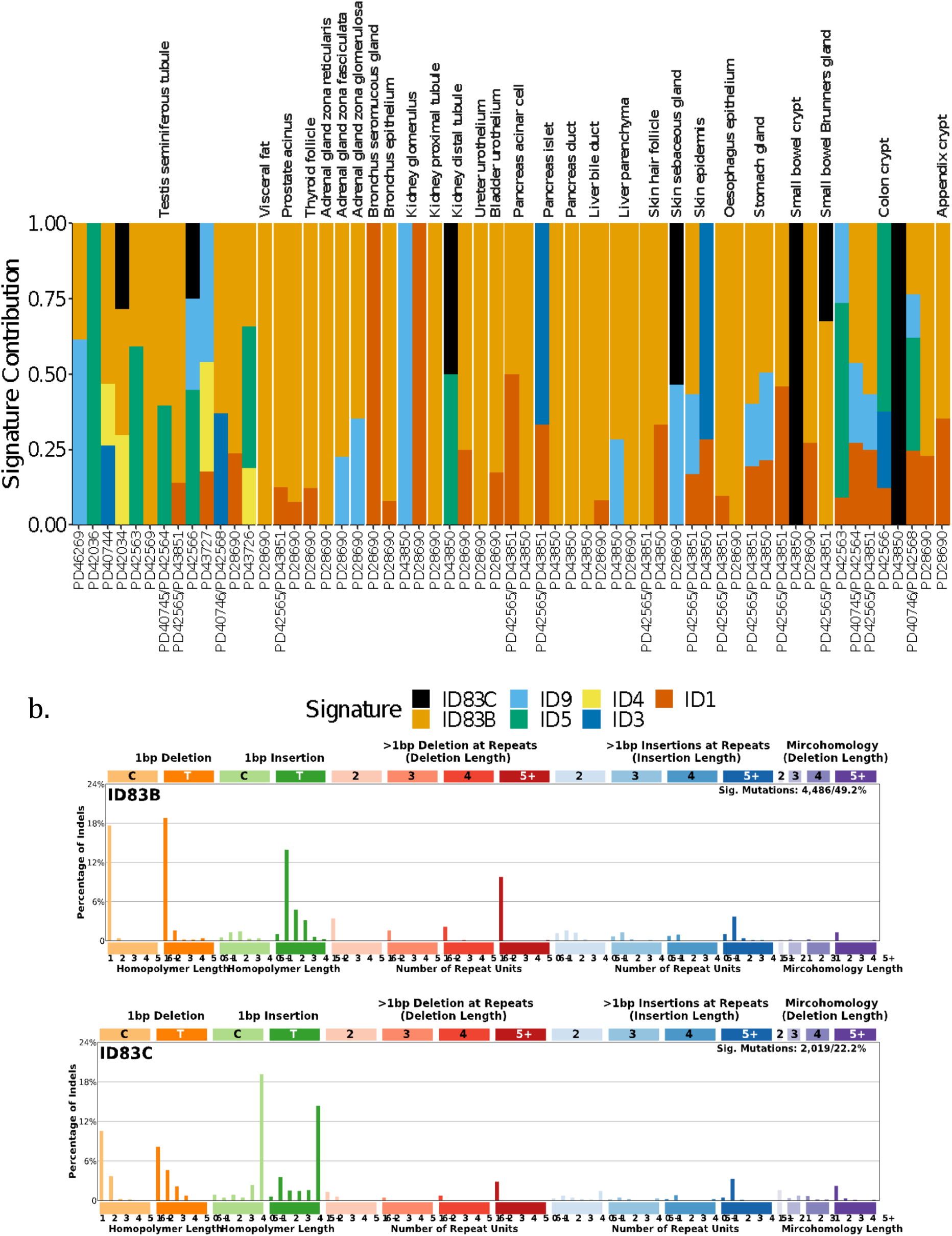
Summary of Indel Signatures: Indels from the same donor were merged together per anatomical structure and signatures were extracted using SigProfiler. **a.** Signature contribution for each donor split by anatomical structure. Donors are sorted by age **b.** the *de novo* signatures extracted by SigProfiler. ID83B contributes almost half of all Indels in the dataset.

**Supplementary Figure 4.**
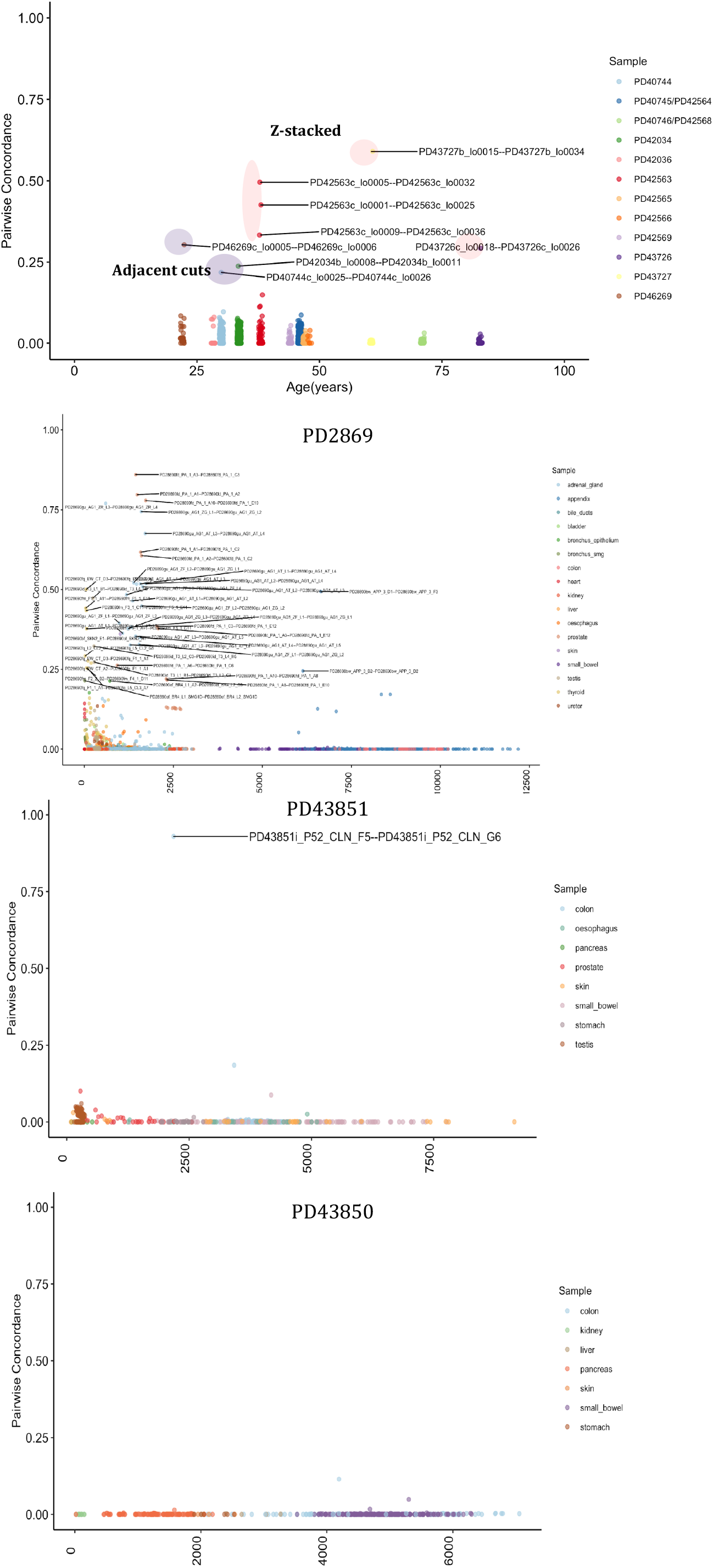
Sample concordance. Pairwise concordance between samples for each donor. Samples sharing a clone or belonging to the same structure are highlighted

**Supplementary Figure 5.**
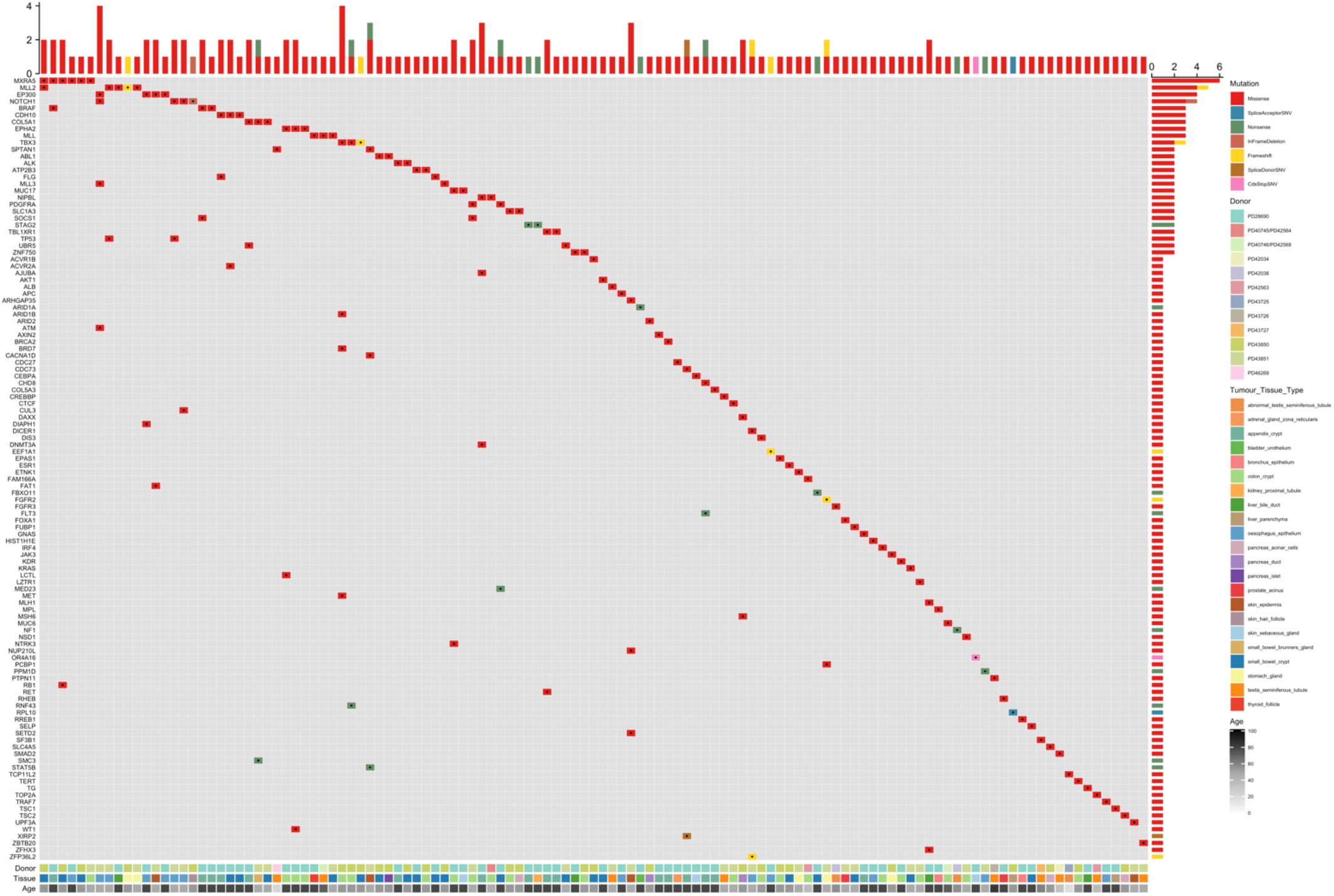
Driver and Passenger mutations in top 100 genes

## Notes

### Competing Interest Statement

The authors have declared no competing interest.

